# Virtual prototyping of non-invasive spinal cord electrical stimulation targeting upper limb motor function

**DOI:** 10.64898/2026.01.22.701010

**Authors:** Abdallah Alashqar, Nabila Brihmat, Vincent Gemar, Zhaoshun Hu, Seong-Ryong Koh, Sandra Diaz-Pier, Roberto de Freitas, Atsushi Sasaki, Matija Milosevic, Rodolfo Keesey, Ismael Seáñez, Esra Neufeld, Hélène Cassoudesalle, Ursula Hofstoetter, Karen Minassian, Fabien Wagner, Andreas Rowald

## Abstract

Transcutaneous spinal cord stimulation (tSCS) applied over the cervical or lumbar spinal cord facilitates motor function after paralysis. However, emerging electrophysiological evidence indicates mechanistic differences between cervical and lumbar tSCS and across different cervical tSCS paradigms. To clarify these discrepancies, we developed and validated a multi-scale, whole-body computational model of tSCS-induced volume conduction, axonal recruitment, and synaptic transmission. Across 24 cervical and four lumbar tSCS paradigms, simulations showed that somatosensory afferents consistently exhibit lower stimulation thresholds than motor efferents. In turn, region-specific synaptic transmission differences may explain electrophysiological discrepancies between cervical and lumbar tSCS. Across cervical tSCS paradigms, substantial volume conduction and axonal recruitment differences were observed that explain electrophysiological discrepancies. Specifically, clinically-prevalent paradigms, including those with anodes placed over the clavicles or iliac crests, engaged peripheral nerves in addition to spinal roots. This effect was amplified by multiphasic waveforms, which introduced recruitment sites near the anodes. By integrating simulations with electrophysiological recordings in 14 able-bodied individuals, we investigated previously unexplored cervical tSCS paradigms on their capacity to recruit somatosensory afferents relative to motor threshold.

## Main

Paralysis, caused by conditions including spinal cord injury (SCI) or stroke, impacts millions worldwide and remains largely without a cure^1^. However, advancements combining epidural spinal cord stimulation (eSCS) with activity-based training have demonstrated remarkable motor function recovery after paralysis^2–9^. Building on this momentum, transcutaneous spinal cord stimulation (tSCS) has been increasingly adopted in clinical research as a non-invasive alternative for motor facilitation^10–21^, driven by the explicit hypothesis that it would emulate the mechanisms of eSCS^19,22,23^. Decades of research support the notion that the motor effects evoked by eSCS are initiated by stimulation-induced action potentials (APs) in somatosensory afferents within the posterior roots^5,24–27,23,28–30^. This afferent-initiated modulation of spinal circuits and their interactions with supraspinal circuits is believed to be essential for promoting neuroplasticity and functional recovery^31–34^. The mechanism of preferential somatosensory afferent over motor efferent recruitment likely translates to lumbar tSCS, as supported by computational modelling^35,36^ and electrophysiological^22,23^ evidence. In turn, clinical results suggest that lumbar tSCS can lead to improvements in weight-bearing and walking after paralysis when combined with activity-based training^11,16–21^.

Recent clinical studies present a similar finding for the upper limbs, demonstrating improvements in hand and arm function when tSCS is applied over the cervical spinal cord^10,12–15,18^. However, the mechanistic framework of eSCS and lumbar tSCS has never been demonstrated to translate to cervical tSCS. Instead, the assumption of preferential somatosensory afferent recruitment with cervical tSCS has recently been challenged by electrophysiological studies^37,38^. These findings highlighted minimal-to-absent post-activation depression (PAD) in paired-pulse experiments around motor threshold (MT), an established electrophysiological metric for assessing the co-activation of somatosensory afferents and motor efferents, suggesting efferent- over afferent-initiated muscle responses^37,38^. Further, muscle-response latencies that match or undershoot peripheral conduction times indicate initiation sites of action potentials (ISAPs) along the anterior-root-peripheral-nerve-axis^37^.

It is important to note that cervical tSCS studies vary widely in the choice of stimulation parameters and not all permutations of stimulation parameters corroborate the same electrophysiological findings^10,12–15,18,37–42^. Among those stimulation parameters, stimulation waveform^37^ and electrode placement^37–39^ have received notable attention in the comparative literature. Recent clinical studies, investigating cervical tSCS for functional rehabilitation, largely employ kilohertz-frequency (KHF) waveforms, with promising clinical outcomes^10,12,13^. Mechanistically, electrophysiological and computational evidence indicates greater co-activation of motor efferents relative to somatosensory afferents with KHF waveforms compared to conventional biphasic pulses^37,43^. Electrode placement also varies substantially across clinical and electrophysiological studies. As multiphasic waveforms involve polarity reversal across pulse phases, electrodes cannot be unambiguously classified as cathodes or anodes. Following established clinical convention^22,23^, we define electrode identity here by polarity during positive pulse phases. Using this convention, clinical studies largely employ paraspinal cathodes and anodes placed over the clavicles or iliac crests^10,12–15,18,37^. These two clinically-prevalent configurations, together with several other anode placements, exhibit minimal-to-absent PAD around MT^37,38^, suggesting that muscle responses are largely initiated by direct recruitment of motor efferents. Only monophasic stimulation with anodes placed over the anterior neck demonstrated an incomplete PAD, suggesting at least a mixed recruitment of somatosensory afferents with motor efferents^38,39,41^. Taken together, these findings suggest that only monophasic pulses applied with neck anodes approximate the mechanisms by which lumbar tSCS elicit muscle responses. In turn, all other cervical tSCS paradigms seemingly engage somatosensory afferents and motor efferents differently than lumbar tSCS.

Here, we hypothesized that, despite paradigm-specific electrophysiological differences, all known biophysical determinants of axonal recruitment^44–47^ indicate that tSCS must universally recruit somatosensory afferents at lower stimulation thresholds than motor efferents. In turn, we speculated that reports of preferential motor efferent recruitment with cervical tSCS may stem from PAD providing an incomplete electrophysiological readout, owing to region-specific synaptic transmission differences^37,48–53^. We posited that variability across cervical paradigms arise from differences in volume conduction and applied waveform, challenging the idea of a single unified cervical tSCS mechanism. To test these hypotheses, we developed a multi-scale computational model of tSCS-induced volume conduction, axonal recruitment, and homonymous monosynaptic transmission across cervical and lumbar tSCS paradigms.

## Results

### Realistic modelling of tSCS

To simulate volume conduction, we developed a 3D model of the human body, including spinal rootlets that extend along peripheral nerves to their innervated muscles (**Fig. 1a,b**). To simulate axonal recruitment, we populated this model with axon trajectories and translated them into conductance-based cable models (**Fig. 1c**), representing populations of somatosensory afferents and motor efferents (**Fig. 1d,e**). For the conductance-based cable model (**Fig. 1c**), we chose the gold-standard in computational modelling of neurostimulation, the McIntyre-Richardson-Grill (MRG) model^54^, as well as a pair of MRG-type models introduced by Gaines et al.^55^, which reproduce key electrophysiological property differences between somatosensory afferents and motor efferents^56,57,47,58^ (**Fig. 1c**). We noted, however, discrepancies in the comparative literature regarding these electrophysiological values, and, in response, tuned another pair of MRG-type models against those previously unmodelled values^44,45^ (**Fig. 1c**, **Extended Data Fig. 1**). To simulate synaptic transmission, we embedded a motoneuron (MN)^32^ model (**Fig. 1f**), and tuned a homonymous monosynaptic transmission model against electrophysiological recordings of peripheral nerve stimulation in the upper and lower limbs^37,48–53^ (**Fig. 1g, Extended Data Fig. 2**).

**Fig. 1.**
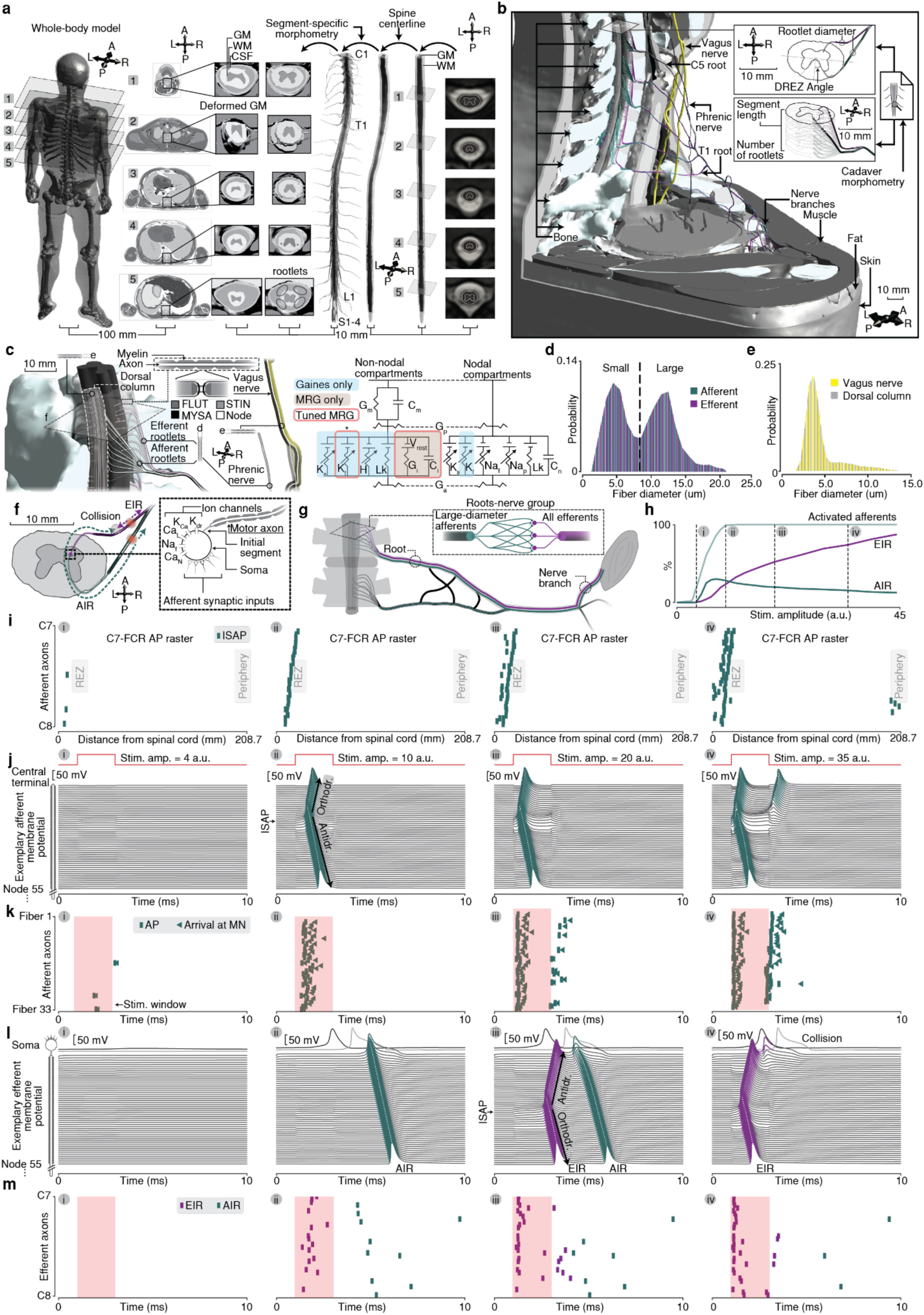
Overview of a multi-scale computational model capable of estimating tSCS-induced volume conduction, axonal recruitment, and homonymous monosynaptic transmission. **a.** 3D model of the entire human body combining an existing whole-body model of a 33-year old adult male composed of 1189 tissues^59^ and a custom-built spinal cord model of an average spinal cord template^60^. **b.** Spinal rootlets were custom-built to fit morphometric data^61–63^ and connected to the peripheral nerves of the whole-body model. **c.** Spinal rootlets, peripheral nerves, and the dorsal column were populated with axon trajectories, which were translated into compartmental cable models^54,55^. Three pairs of compartmental cable models were implemented: The MRG-model^54^, the MRG-type model pair implemented by Gaines et al.^55^, and a newly-tuned MRG-type model pair (see Extended Data Fig. 1). Highlighted parts refer to unique ion channels in each of the models. *The fast potassium channel included in the newly-tuned MRG-type model pair was only added to the fluted (FLUT) internodal segments, while it was present in all non-nodal segments of the MRG-type model pair of Gaines et al.^55^. **d.** Properties of the compartmental cable models were scaled by axon diameter to represent axon populations. Depicted are the axon diameter distributions used for the cervical somatosensory afferents and motor efferents in the spinal rootlets^46^. **e.** Similarly to (d) properties of compartmental cable models in the vagus nerve and dorsal column were scaled by axon diameter. Depicted are the axon diameter distributions used for the dorsal column and vagus nerve axons^64^. **f.** A homonymous monosynaptic transmission model (see Extended Data Fig. 2) was implemented by projecting the compartmental cable models representing somatosensory afferents to motoneuron (MN) models, themselves connected to compartmental cable models representing motor efferents. Action potentials (APs) in the motor efferents induced transsynaptically by somatosensory afferents were defined as afferent-initiated responses (AIRs), whereas direct stimulation-induced APs in the motor efferents were defined as efferent-initiated responses (EIRs). **g.** Synaptic transmission properties including projections between somatosensory afferents and motor efferents were based on physiological reports^49–53,65^, as well as tuned against peripheral nerve stimulation experiments^37,48^ (see Extended Data Fig. 2). **h.** An exemplary recruitment curve of somatosensory afferents, AIR, and EIR is shown. Notably, multiple somatosensory afferents can be recruited at a stimulation intensity that is insufficient to elicit an AIR (i). With increasing stimulation amplitude, AIR- and EIR-curves exhibit the characteristic shapes of H-reflexes and M-waves^48^ (ii, iii, iv). **i.** Exemplary initiation sites of APs in somatosensory afferents are depicted for each stimulation intensity i-iv from (**h**). **j.** The nodal membrane potentials of an exemplary somatosensory afferent are depicted for each stimulation intensity i-iv from (**h**). **k.** Exemplary temporal pattern of APs in somatosensory afferents and arrival at MNs are depicted for each stimulation intensity i-iv from (**h**). **l.** The nodal membrane potentials of an exemplary motor efferent are depicted for each stimulation intensity i-iv from (**h**). **m.** Exemplary temporal pattern of APs in motor efferents are depicted for each stimulation intensity i-iv from (**h**). Abbreviations: anterior (A), posterior (P), right (R), left (L), gray matter (GM), white matter (WM), cerebrospinal fluid (CSF), dorsal root entry zone (DREZ), millimeter (mm), fluted (FLUT), stereotyped internode (STIN), myelin sheath attachment (MYSA), myelin conductance (G_m_), myelin capacitance (C_m_), periaxonal conductance (G_p_), axoplasmic conductance (G_a_), slow potassium (K_s_), fast potassium (K_f_), hyperpolarization-activated cyclic-nucleotide-gated (HCN), leakage (Lk), internodal conductance (G_i_), internodal capacitance (C_i_), resting membrane potential (V_rest_), persistent sodium (Na_p_), fast sodium (Na_f_), nodal capacitance (C_n_), N-type calcium (Ca_N_), L-type calcium (Ca_L_), calcium-activated potassium (K_Ca_), rectifier potassium (K_dr_), flexor capri radialis (FCR), root entry zone (REZ), stimulus (stim.), amplitude (amp.), micrometer (μm), arbitrary units (a.u.), millivolt (mV).

By combining all these components, our model provides a virtual prototyping platform for neurostimulation paradigms, enabling the simulation of the initiation and propagation of APs in somatosensory afferents (**Fig. 1i,j,k**) and motor efferents (**Fig. 1l,m**), as well as the transsynaptic induction of afferent-initiated responses (AIR) (**Fig. 1h,l**), relative to efferent-initiated responses (EIR) (**Fig. 1h,l,m**) as proxies for muscle responses. We then validated the accuracy of our model against electrophysiological datasets from six cervical and one lumbar tSCS paradigms^37,38^ (**Fig. 2, Extended Data Fig. 3**). First, we estimated MT as the lowest stimulation amplitude required to elicit either an AIR or an EIR and found that our newly introduced MRG-type model pair best reproduced MT trends across muscles (**Fig. 2b-g**). Second, we reasoned that paradigm-dependent differences in PAD offer an experimental analogue to relative differences in the percentage of AIRs (PAIRs) in our model (**Fig. 2a**). Comparing the difference between trends of normalized PAD and PAIR indicated that our newly-tuned MRG-type model pair closely accounts for recruitment differences across tSCS paradigms (**Fig. 2h,i, Extended Data Fig. 3e,f**).

**Fig. 2.**
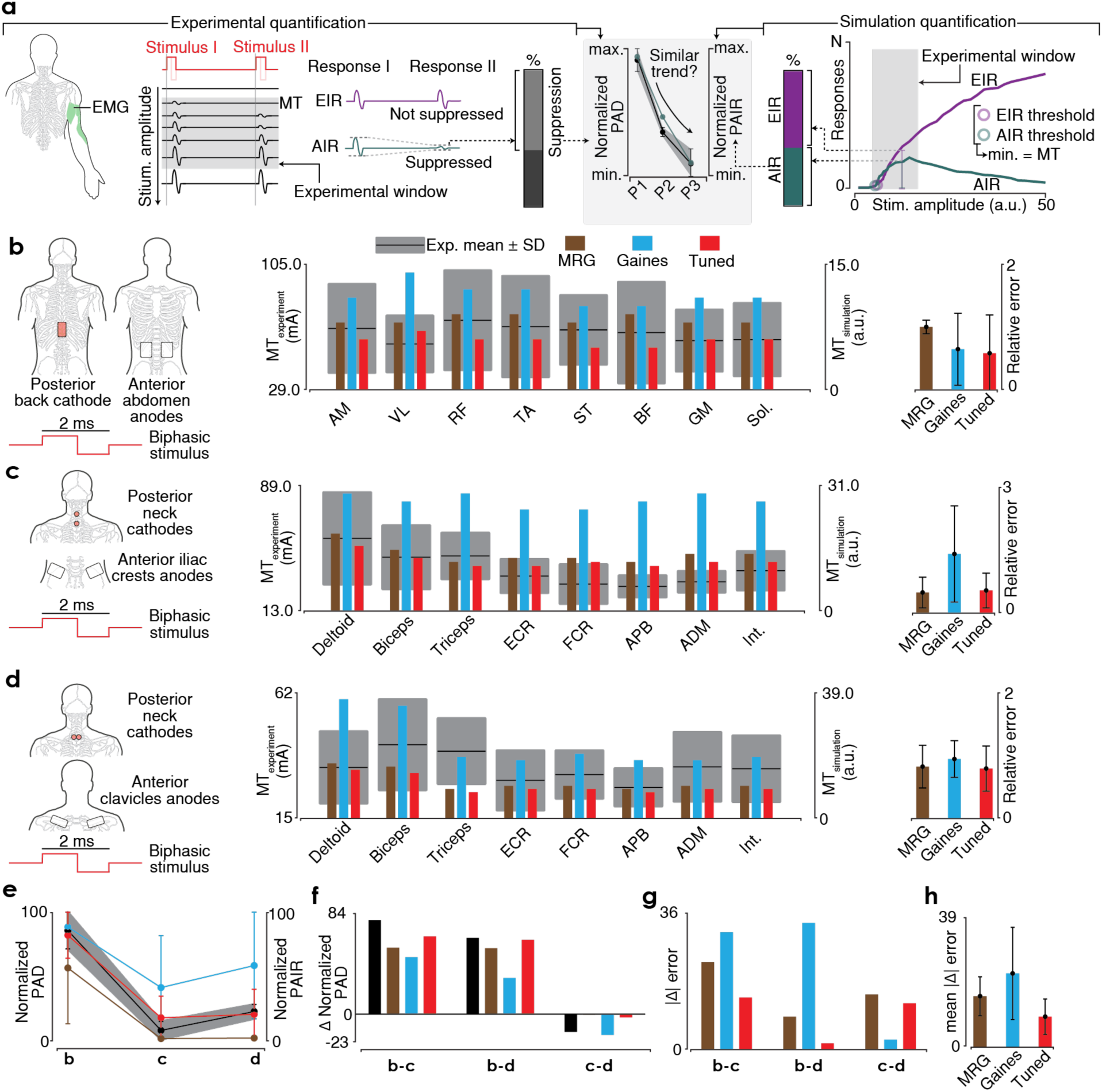
Validation of the computational model against previously reported electrophysiological recordings of cervical and lumbar tSCS^37^. **a.** We validated our computational model through a two-step process. First, we reasoned that the afferent- or efferent-initiated response (AIR or EIR) elicited at the lowest possible stimulation amplitude provides a computational approximation of motor threshold (MT) within an arbitrary electrophysiological tSCS experiment (**b-d**). Second, we aimed to validate our model’s capacity to accurately calculate the relative co-activation of somatosensory afferents and motor efferents, by introducing the percentage of AIRs (PAIR) to quantify the recruitment of somatosensory afferents relative to motor efferents. This relative co-activation of somatosensory afferents and motor efferents is commonly extrapolated from post-activation depression (PAD) obtained with paired-pulse experiments. However, PAD is not a pure expression of this relative co-activation^66^. In turn, we reasoned that differences between PAD values derived from different tSCS paradigms within the same study population and with the same paired-pulse protocol must stem from differences in the relative co-activation of somatosensory afferents and motor efferents. We thus validated our computational model by comparing trends of PAD and PAIR across tSCS paradigms (**e-h**). As part of the validation process the compartmental cable models were systematically varied between the three model pairs introduced in Fig. 1c to assess their influence on MT and the relative co-activation of somatosensory afferents and motor efferents. **b-d.** Each row depicts a sketch of the applied tSCS paradigms (left), the comparison between experimental and simulated MT across different muscles (middle), and the relative error between the normalized experimental and simulated MT (right). **e**. Comparison of normalized PAD (black line represents the average value whereas the gray area represents the standard deviation) and PAIR (colored lines) across the different tSCS paradigms from (**b-d**). **f**. Differences in PAD between tSCS paradigms (black bar) and differences in PAIR between tSCS paradigms (colored bars). **g**. Differences between PAD and PAIR across trends comparing tSCS paradigms. **h.** Average and standard deviation of (**g**) across trends of tSCS paradigms. Abbreviations: electromyography (EMG), paradigm (P), minimum (min.), maximum (max.), stimulus (stim.), millisecond (ms), milliampere (mA) adductor magnus (AM), vastus lateralis (VL), rectus femoris (RF), tibialis anterior (TA), semitendinosus (ST), biceps femoris (BF), gastrocnemius (GM), soleus (Sol.), experiment (Exp.), standard deviation (SD) extensor carpi radialis (ECR), flexor carpi radialis (FCR), abductor pollicis brevis (APB), abductor digiti minimi (ADM), interosseous (Int.).

### tSCS preferentially recruits somatosensory afferents

We next aimed to elucidate the differential recruitment mechanisms across tSCS paradigms using our validated computational model (**Fig. 3, 4, Extended Data Fig. 4-7**). We calculated axonal recruitment curves and derived AIR- and EIR-thresholds across six cervical and one lumbar electrode placements with monophasic, biphasic and KHF waveforms (**Fig. 3, Extended Data Fig. 4,5**). We found that, despite paradigm-specific differences, somatosensory afferents were recruited at lower stimulation thresholds than motor efferents for all tSCS paradigms (**Fig. 3, Extended Data Fig. 4,5**). Importantly, the AIR-threshold depends on the synaptic transmission model, which differs between cervical and lumbar segments^37,48–53^ (**Extended Data Fig. 2**). Taking this element into account, we found that EIR-thresholds were lower than AIR-thresholds for all cervical tSCS paradigms other than monophasic stimulation with a neck anode, for which AIR and EIR were identical (**Fig. 3, Extended Data Fig. 4**). By contrast, the EIR-threshold was larger than the AIR-threshold for lumbar tSCS (**Extended Data Fig. 5a-d**). These simulated findings align with previous electrophysiological reports on PAD differences across cervical and lumbar tSCS paradigms^37,38^. We thus concluded that although all tSCS paradigms predominantly recruit somatosensory afferents at clinically-relevant sub-MT stimulation amplitudes, muscle responses at MT are elicited through direct motor efferent recruitment in all cervical tSCS paradigms other than monophasic stimulation with a neck anode, for which both afferent- and efferent-mediated mechanisms elicit muscle responses at MT (**Fig. 3, Extended Data Fig. 4,5**).

**Fig. 3.**
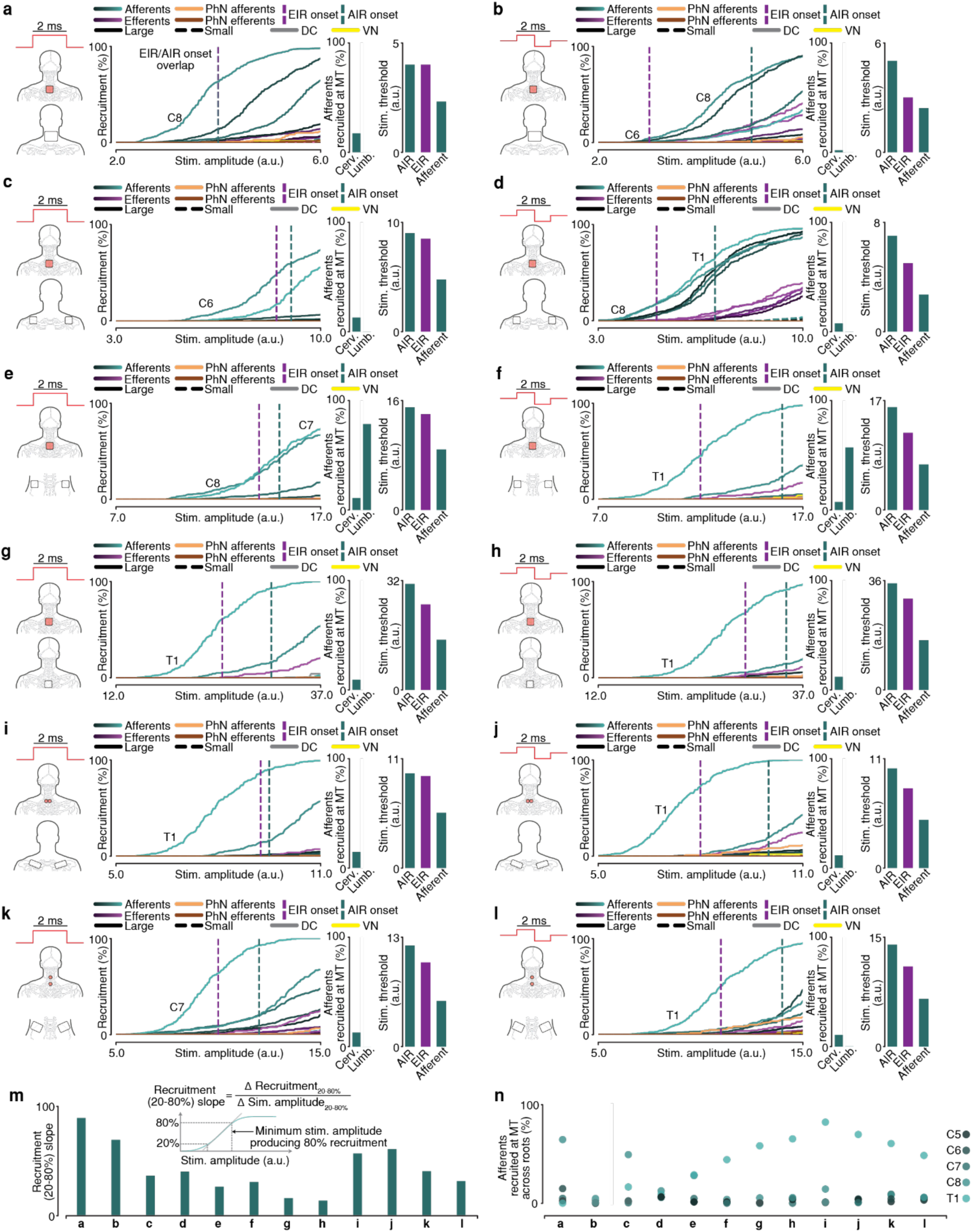
All common cervical tSCS paradigms^37,38^ exhibit lower stimulation thresholds for somatosensory afferents than motor efferents in-silico. However, efferent-initiated responses dominate motor threshold (MT) in all but monophasic neck-anode stimulation. **a-l** From left to right: Stimulation configurations, recruitment curves of all simulated axons including afferent-initiated (AIR) and efferent-initiated response (EIR) thresholds, percentage of somatosensory afferents recruited at MT, and stimulation thresholds for AIRs, EIRs, and afferent recruitment (right) for six commonly used cathode and anode placements. Monophasic stimulation is shown in (**a,c,e,g,i,k**), and biphasic stimulation in (**b,d,f,h,j,l**). For inverted monophasic polarity and kilohertz-frequency waveforms please refer to Extended Data Fig. 4. **m.** Recruitment slopes of somatosensory afferents across all cervical tSCS paradigms (**a–l**), are shown as an indicator of variance in the efficacy of somatosensory afferent recruitment relative to stimulation amplitude. **n**. Somatosensory afferents recruited at MT across posterior roots for all cervical tSCS paradigms (**a-l**) are depicted as an indicator of variance in the recruitment of somatosensory afferents between spinal roots. Abbreviations: millisecond (ms), phrenic nerve (PhN), dorsal column (DC), vagus nerve (VN), cervical (Cerv.), lumbar (Lumb.), motor threshold (MT), stimulus (stim.), arbitrary units (a.u.)

### Cervical tSCS recruits peripheral nerves

We next aimed to reconcile the mechanistic ambiguities between different cervical tSCS paradigms. To this end, we visualized the volume-conduction differences and the second spatial derivative of the extracellular potential along the axons (∂^2^V/∂X^2^)^67^, which constitutes a proxy for stimulus-induced axonal recruitment at its local maxima (max. ∂^2^V/∂X^2^)^67^ (**Fig. 4, Extended Data Fig. 5,7**). Cervical tSCS with anodes over the anterior neck directs current through the midline of the body (**Fig. 4a**), resulting in max. ∂^2^V/∂X^2^ in the spinal roots (**Fig. 4a**), thus concentrating ISAPs in the spinal roots (**Fig. 4b**). Similarly, lumbar tSCS with anodes over the abdomen (**Extended Data Fig. 5a-d**) directs current through the spine (**Extended Data Fig. 5e**), resulting in max. ∂^2^V/∂X^2^ and ISAPs in the spinal roots (**Extended Data Fig. 5f-h**). In contrast, all other anode positions, including the clinically-prevalent positions over the clavicles (**Fig. 4c,d**) or iliac crests (**Fig. 4e,f**), direct current towards the periphery (**Fig. 4c,e**), adding max. ∂^2^V/∂X^2^ and ISAPs in peripheral nerves (**Fig. 4d,f)**. In peripheral nerves, somatosensory afferents and motor efferents are not spatially segregated and therefore experience near-identical extracellular voltage. In contrast, posterior and anterior roots are spatially separated and exposed to different voltages. Consequently, tSCS paradigms that place the anodes distal to the neck (hereafter collectively referred to as distal tSCS) produce greater motor-efferent co-activation than cervical tSCS with anterior neck anodes (hereafter referred to as neck tSCS) (**Fig. 3, Extended Data Fig. 4**).

**Fig. 4.**
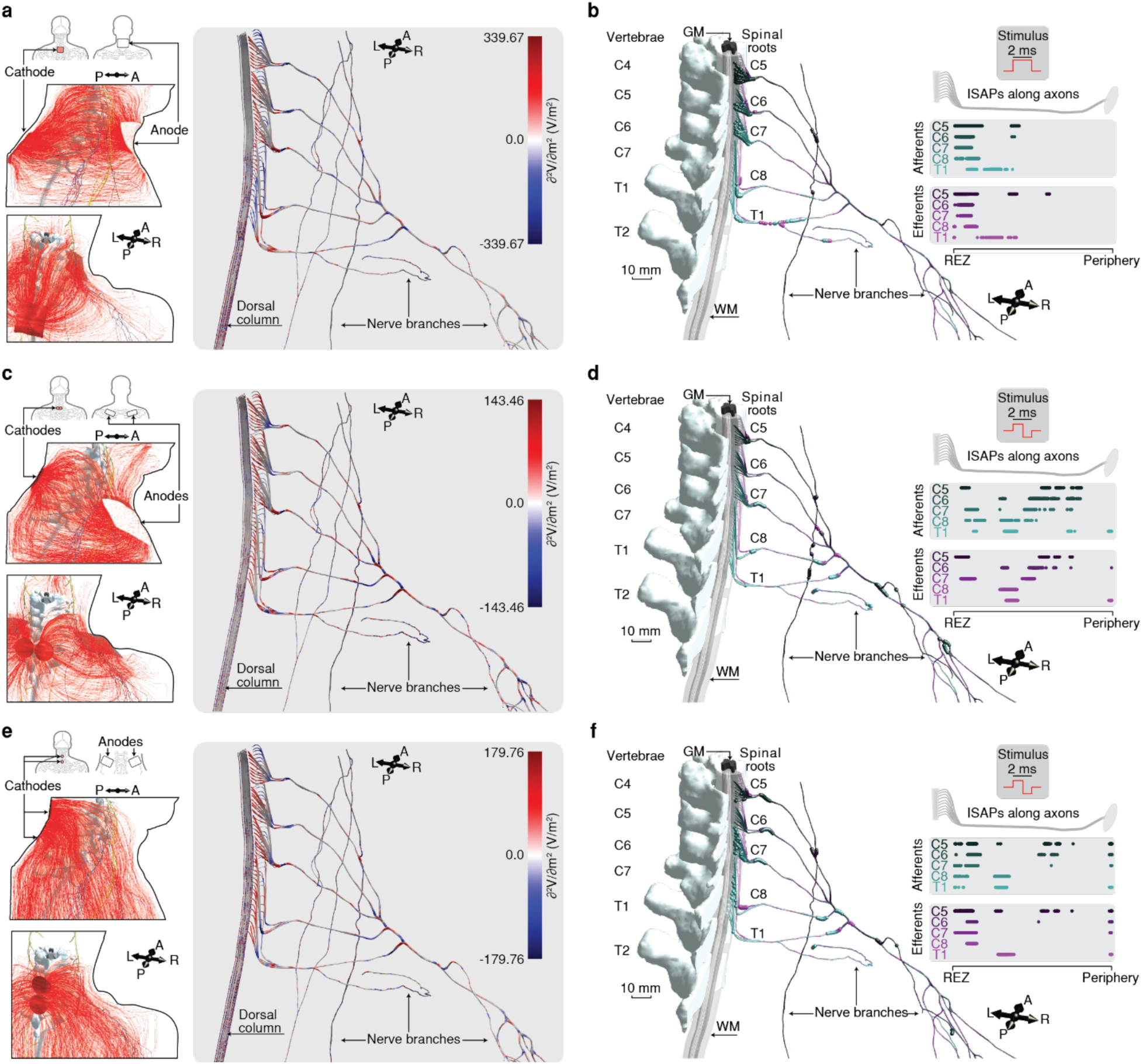
Simulations indicate that neck tSCS directs current through the spine, thus concentrating initiation sites of action potentials (ISAPs) in the spinal rootlets. In turn, distal tSCS directs current towards the periphery, adding ISAPs in peripheral nerves. **a,c,e.** Electric field streamlines (left) and the corresponding second spatial derivative of the extracellular potential along the axons (∂^2^V/∂X^2^) governing neural recruitment (right) for three commonly used cathode–anode placements^37,38^. **b,d,f.** ISAP distributions for somatosensory afferents (turquoise) and motor efferents (purple) corresponding to the stimulation paradigms shown in (**a,c,e**). Abbreviations: anterior (A), posterior (P), right (R), left (L), millimeter (mm), grey matter (GM), white matter (WM), root entry zone (REZ), initiation sites of action potentials (ISAPs), volt (V), meter (m).

We next reasoned that this peripheral-nerve-recruitment mechanism must be impacted by the negative phases of multiphasic waveforms. We compared monophasic pulses of either polarity with a biphasic waveform for neck **(Extended Data Fig. 6a)** and distal tSCS **(Extended Data Fig. 6b)**. For neck tSCS, biphasic ISAPs approximate the superposition of monophasic pulses of either polarity (**Extended Data Fig. 6a**), for which the positive phase favored posterior-root somatosensory afferents, whereas the negative phase favored anterior-root motor efferents. In turn, for distal tSCS, the ISAPs during the biphasic pulse resemble the ISAPs during the anodic monophasic pulse, both concentrating ISAPs in the peripheral nerves, whereas cathodic monophasic stimulation distributes ISAPs along the entire spinal-root–peripheral-nerve-axis (**Extended Data Fig. 6b**). In the case of iliac crest anodes, the current is directed through the entire upper body (**Extended Data Fig. 7a,d**), resulting in max. ∂^2^V/∂X^2^ and ISAPs in both cervical and lumbar structures (**Extended Data Fig. 7b,c,e,f**), which may result in co-activation of lumbar relative to cervical axons (**Fig. 3e,f,k,l**). This peripheral-nerve-recruitment mechanism elicited by distal tSCS and amplified by multiphasic waveforms explains both the PAD differences reported across cervical tSCS paradigms^38^ and the short motor-response latencies observed with distal anodes^37^, whereas neck tSCS predominantly recruits axons within the spinal roots.

### Anode size and position govern axonal recruitment

Importantly, somatosensory afferent recruitment varied widely across cervical tSCS paradigms (**Fig. 3m,n, Extended Data Fig. 4m,n)**, suggesting the existence of a paradigm that maximizes somatosensory afferent recruitment at clinically-relevant sub-MT amplitudes (**Fig. 5, Extended Data Fig. 8-10**). Based on our novel mechanistic framework, maximal somatosensory afferent recruitment is achieved with neck tSCS (**Fig. 3a,b)**. However, this paradigm may pose feasibility challenges^37^ in clinical translation studies, indicating a need for identifying an alternative anode placement. We thus investigated seven anode placements (**Fig. 5a**), four anode sizes (**Fig. 5d**), three pulse widths (**Fig. 5g**), nine cathode placements (**Fig. 5j**), and the impact of vertebral vs intervertebral cathode positioning (**Fig. 5m**).

**Fig. 5:**
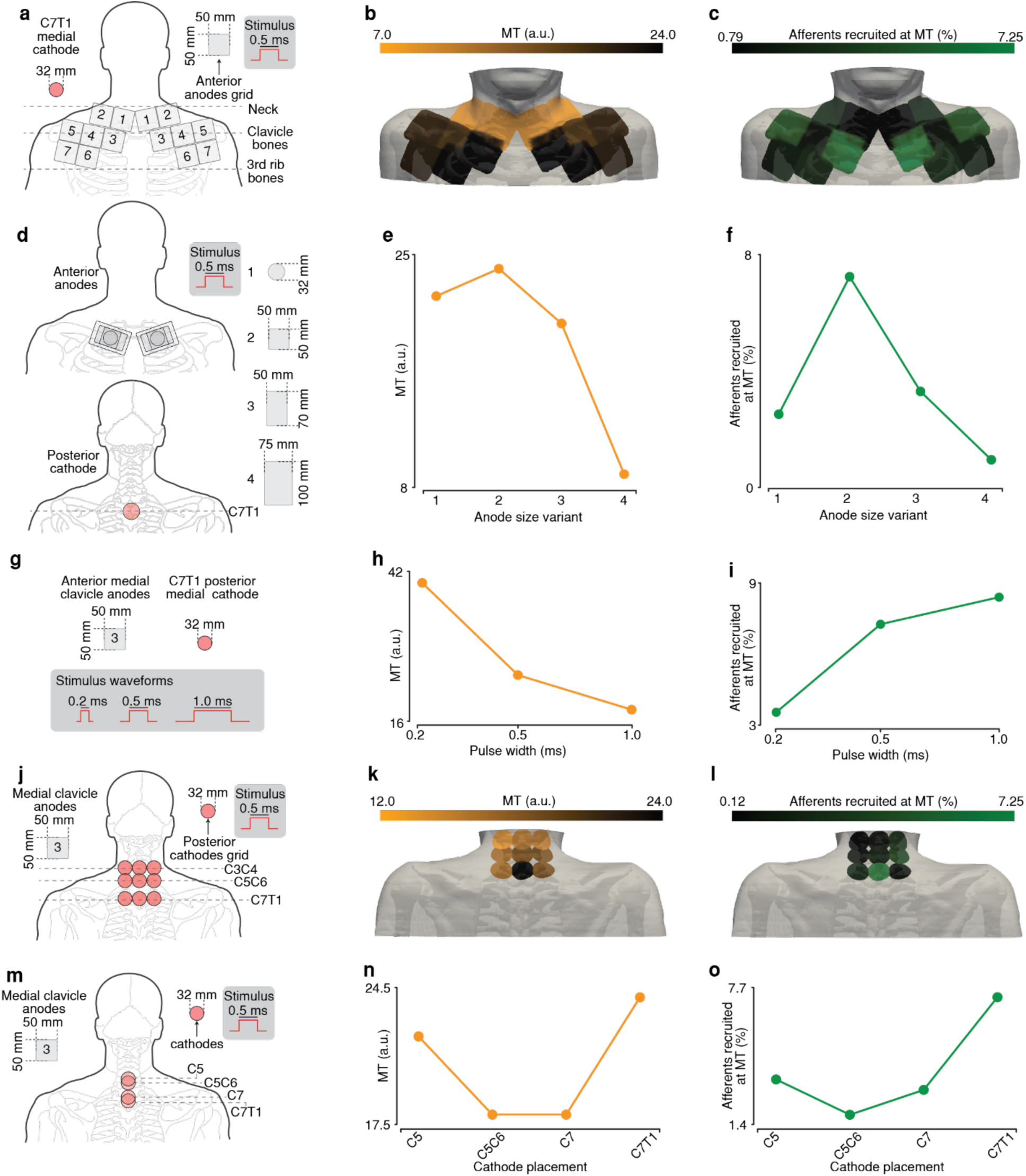
50x50mm anodes placed over the clavicles proximal to the neck maximize somatosensory afferent recruitment, whereas mid-clavicular placement and larger anodes lowered motor threshold (MT). **a,d,g,j,m.** Sketch of tSCS parameters used for exploring the impact of anodal placement (**a-c**), anodal size (**d-f**), pulse width (**g-i**), cathodal placement (**j-l**), and vertebral vs. intervertebral cathode position (**m-o**) on MT and somatosensory afferent recruitment. **b,e,h,k,n.** MT for each tSCS parameter of (**a,d,g,j,m**). **c,f,I,l,o.** Percentage of somatosensory afferents recruited at MT for each tSCS parameter of (**a,d,g,j,m**). Abbreviations: millimeter (mm), arbitrary units (a.u.), millisecond (ms).

Our simulations indicated that MT increases with increasing distance of the anode from the neck (**Fig. 5b**), with C5 and C6 roots as well as associated deltoid and biceps muscles expressing greater variation in MT than other roots and muscles in our model (**Extended Data Fig. 10a**). Recruitment of somatosensory afferents is maximized with anodes proximal to the neck (**Fig. 5c**). MT and somatosensory afferent recruitment exhibited similar trends with size variations of the anodes, with 50x50 mm anodes maximizing both MT and somatosensory afferent recruitment (**Fig. 5e,f**). Increasing pulse width decreased MT (**Fig. 5h**) and increased somatosensory afferent recruitment (**Fig. 5i)**. Alterations to cathode positioning did not substantially change MT, except for the most caudally placed cathode along the midline (C7T1), which maximized MT (**Fig. 5k, Extended Data Fig. 10b**). In contrast, this midline-C7T1 cathode position maximized recruitment of somatosensory afferents (**Fig. 5l**). No clear trend in cathode positioning over vertebral vs intervertebral space was observed (**Fig. 5n,o**). We further noticed substantial left-right-asymmetries across cathode positions (**Fig. 5k,l, Extended Data Fig. 10b**), arising due to left-right-asymmetries of nerve trajectories in our model (**Extended Data Fig. 9**). Potential co-activation of other nerves unrelated to the C5-T1 spinal roots remained minimal across all stimulation parameters (**Extended Data Fig. 8**). These results motivate the clavicular placement of 50x50mm anodes proximal to the neck to maximize somatosensory afferent recruitment, while mid-clavicular placement and larger anodes may reduce MT.

Finally, we electrophysiologically validated these predictions in 14 able-bodied individuals. We reduced the parameter space to three anode positions covering the clavicles (**Fig. 6a**), two cathode positions (**Fig. 6d**), three anode sizes (**Fig. 6g**), and three pulse widths (**Fig. 6j**). Importantly, direct recordings of somatosensory afferent recruitment pose obvious feasibility issues during electrophysiological experiments in humans, while our simulations suggested that the paired-pulse method would yield minimal-to-absent PAD at MT without revealing detailed information on axonal recruitment. Indeed, PAD remained minimal-to-absent across all tested tSCS parameters (**Fig. 6c,f,i,l**). In turn, we leveraged MT to confirm our simulations. MT exhibited a statistically significant minimum at the mid-clavicular position (p < 0.05) (**Fig. 6b**). MT increased with caudal displacement of cathodes (p < 0.05) (**Fig. 6e**). No statistically significant change was observed between 32 mm and 50x50 mm anodes, yet 50x70 mm anodes exhibited statistically significant decreases in MT relative to smaller anodes (p < 0.05) (**Fig. 6h**). MT decreased statistically significantly with increasing pulse width (p < 0.05) (**Fig. 6k**).

**Fig. 6.**
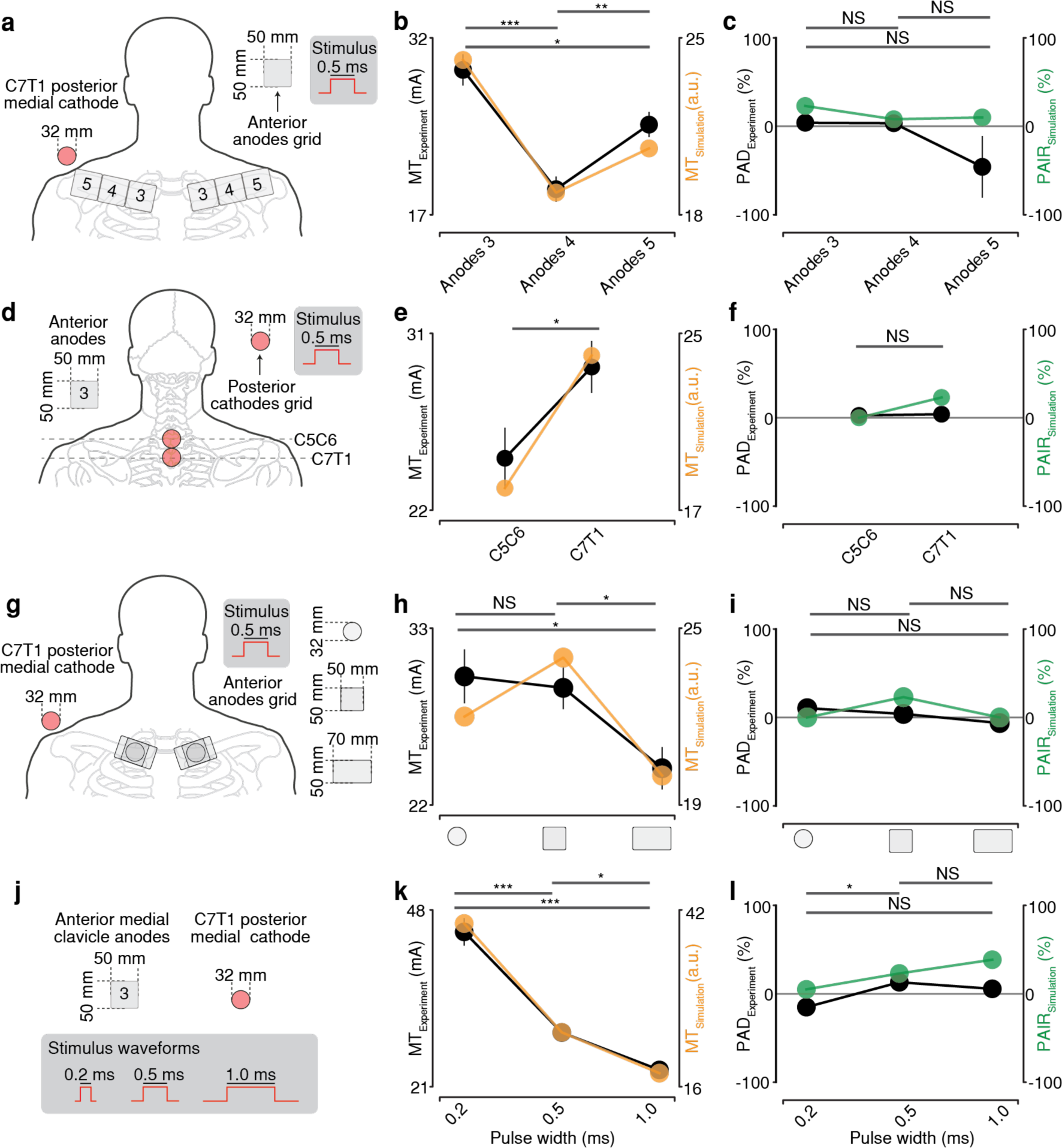
Validation of simulated trends against electrophysiological recordings in able-bodied individuals (n=14). **a,d,g,j.** Sketch of tSCS parameters used for comparing simulations against experiments. **b,e,h,k.** Motor threshold of simulation (orange) and experiment (black) for each tSCS parameter of (**a,d,g,j**). **c,f,i,l.** Post-activation depression (black) and percentage of afferent-initiated responses (green) for each tSCS parameter of (**a,d,g,j**). Black error bars represent the standard error of the mean (SEM). Statistics represent one-way ANOVA with Bonferroni post-hoc correction. *P < 0.05, **P < 0.01 and ***P < 0.001. NS, not significant. Abbreviations: millimeter (mm), millisecond (ms), motor threshold (MT), post-activation depression (PAD), percentage of afferent-initiated responses (PAIR).

## Discussion

### Novel findings in tSCS

The original goal of tSCS was to provide a non-invasive alternative to eSCS by selectively activating somatosensory afferents in the posterior roots to modulate spinal circuits^19,22^. Yet, these biophysical foundations have never been demonstrated to translate from lumbar to cervical tSCS. A previous computational study calculated lower stimulation thresholds of somatosensory afferents than motor efferents with cervical tSCS, however simulated only neck tSCS with monophasic pulses in a simplified volume conductor model^68^. While electrophysiological recordings with neck tSCS suggest that muscle responses are at least partially initiated by somatosensory afferents^38,39,41^, recent electrophysiological findings of distal tSCS with multiphasic waveforms, report minimal-to-absent PAD during paired-pulse experiments and motor response latencies equal to or shorter than peripheral conduction times^37^. Our computational results reconcile these discrepancies between cervical and lumbar tSCS, as well as among different cervical tSCS paradigms through three mechanistic insights.

First, weaker monosynaptic transmission in the cervical compared to the lumbar spinal cord (**Extended Data Fig. 2**) necessitates a greater contribution of somatosensory afferents to initiate AIRs during cervical tSCS. This requirement elevated AIR-thresholds relative to EIR-thresholds (**Fig. 3, Extended Data Fig. 4,5**), thus providing a mechanistic explanation for the reduced PAD across cervical tSCS paradigms compared to lumbar tSCS^37,38^. Importantly, these region-specific synaptic transmission differences imply that PAD systematically underestimates the engagement of somatosensory afferents at MT in cervical tSCS.

Second, while neck tSCS directs current through the spine (**Fig. 4a**), distal tSCS directs current towards the periphery (**Fig. 4c,e**). These volume conduction differences result in preferential recruitment at spinal roots for neck tSCS (**Fig. 4b, Extended Data Fig. 6a**), while distal tSCS additionally recruits peripheral nerves (**Fig. 4d,f**, **Extended Data Fig. 6b,7**). While somatosensory afferents and motor efferents are housed in spatially separated posterior and anterior roots respectively, they converge in the peripheral nerves. Thus, total electric field strength differences between posterior and anterior roots result in increased somatosensory afferent recruitment relative to motor efferents for neck tSCS compared to distal tSCS with monophasic pulses (**Fig. 3a,c,e,g,i,k,m,n**).

Third, during multiphasic waveforms, cathodal and anodal stimulation invert at polarity reversal, causing recruitment at different ISAPs. For neck tSCS, these negative phases favor anterior-root motor efferent recruitment (**Fig. 3b, Extended Data Fig. 4 a,b**). For distal tSCS, the negative phases concentrate ISAPs within peripheral nerves (**Extended Data Fig. 6b,7**), further shifting recruitment mechanisms of tSCS away from eSCS-like spinal root recruitment toward peripheral nerve stimulation, and elevating co-activation of motor efferents relative to somatosensory afferents (**Fig. 3d,f,h,j,l,m,n, Extended Data Fig. 4c-n**).

Together, these anode- and phase-dependent peripheral-nerve-vs-spinal-root recruitment differences explain the smaller PAD observed with distal tSCS compared with neck tSCS^38^ and motor response latencies equal or shorter to peripheral conduction times^37,38^. The implications of directing current through the entire body with distal tSCS even extend to recruitment of lumbar axons with iliac crest anodes (**Fig. 3e, Extended Data Fig. 4e,7**), providing a possible explanation for observations of lower limb motor facilitation with this tSCS paradigm^42^. Notably, we did not explicitly calculate the recruitment of other neural substrates like thoracic roots and associated nerves, which may also be engaged with cervical tSCS paradigms that direct current through large volumes of the body.

Importantly, our simulations indicate substantial differences in volume conduction and axonal recruitment across cervical tSCS paradigms (**Fig. 3,4, Extended Data Fig. 4-7**). Given these complex biophysical mechanisms of cervical tSCS and the lack of comparative electrophysiological analysis, there is little consensus on effective stimulation parameters^10,12–15,18,37–42^. A previous computational study of cervical tSCS ventured into simulation-based optimization of stimulation parameters, focusing on cathode placement along the midline of the posterior spinal cord and cathode size^69^. Given our mechanistic insights into the importance of anodes, we focused our optimization on their position and size (**Fig. 5,6, Extended Data Fig. 8,10**). We identified that 50x50 mm anodes placed over the clavicles proximal to the neck maximizes somatosensory afferent recruitment, whereas mid-clavicular placement or larger sizes reduce MT (**Fig. 5**,**6**). We provided electrophysiological recordings of 14 able-bodied individuals that corroborate our computational findings (**Fig. 6**). Future studies ought to evaluate the clinical efficacy of the identified stimulation parameters in individuals with paralysis. Finally, volume conduction asymmetries observed in our model (**Extended Data Fig. 9**) suggest that anatomical variability should be considered when applying the proposed stimulation parameters.

### Novel findings in computational modelling

Computational models, combining volume conductor with compartmental cable models, are widely used to virtually prototype spinal cord neurostimulation^5,24,32,35,36,68,70^. While such models offer valuable translational insights^5,24,32,35,36,68,70^, key methodological questions remain unresolved^70^.

One open question concerns the required level of anatomical realism in volume conductor models^71,72^. This may be especially pertinent for non-invasive paradigms like tSCS, where current traverses multiple tissue layers to reach neural targets. Indeed, we demonstrate here how inclusion of peripheral nerves uncovered peripheral-nerve-recruitment mechanisms with distal tSCS and substantial volume conduction and axonal recruitment differences across tSCS paradigms (**Fig. 3,4, Extended Data Fig. 4-7,9**). These mechanisms could not have been detected with contemporary models that focus on spinal structures^35,68^, highlighting an inherent benefit to virtually prototype neurostimulation in anatomically realistic environments. Our model may be understood as a contemporary maximally realistic solution, representing over 1189 tissues of the human body^59^ (**Fig. 1a**), and integrating multi-modal data^46,60–64^ to represent spinal nerves and rootlets, capturing their trajectories^63^, bending angles^62^, orientations^63^, entry zones^46,64^, and diameter distributions^46,64^ (**Fig. 1b,d,e**). This level of realism necessitated extraordinary computational resources (see Methods), highlighting both the increasingly important role of high-performance computing in computational neuroscience^73^ and the need for future investigations to determine minimally viable solutions of anatomical realism.

Another critical question remains in the accuracy of conductance-based cable models. The MRG-model has often been used to represent a broad range of axons, including somatosensory afferents^36,54,74^, despite not explicitly representing their specific electrophysiological properties (**Extended Data Fig. 1d**). A previous study by Gaines et al. has addressed this limitation by introducing a pair of sensory and motor variants^55^. While the Gaines-model-pair successfully reproduces key electrophysiological differences between somatosensory afferents and motor efferents^55^, the underlying values vary between sources^44,47^. In response, we introduced a novel MRG-type pair tuned against previously unmodelled values^44,45^ (**Fig. 1c, Extended Data Fig. 1**). In comparison to the MRG-model^54^ and the Gaines-model-pair^55^, our newly-tuned MRG-type pair performed best in our validation pipeline (**Fig. 2, Extended Data Fig. 3**). We emphasize however, remaining uncertainties in the simulation of axonal recruitment, arising, for instance, from variability in estimates of somatosensory afferent and motor efferent diameters, which propagate into population-level compartmental cable model predictions^75–78^. Extending the introduced tuning pipeline to sub-populations of somatosensory afferents (e.g. proprioceptive and cutaneous) may enable finer mechanistic dissection across neurostimulation modalities.

A critical question across computational investigations of neurostimulation is how to operationalize simulated findings into experimental settings. Contemporary computational models provide insights into the interactions between neurostimulation and neural substrates yet fail to connect to electrophysiological studies providing insights into what is observed at the muscles. A key component introduced here is a homonymous monosynaptic transmission model^49,50,53^ (**Fig. 1f-m, Extended Data Fig. 2**), enabling direct comparison with electrophysiological observations (**Fig. 2,6, Extended Data Fig. 2,3**). We tuned our model against peripheral nerve recruitment experiments (**Extended Data Fig. 2**), extensively validated the capacity of our model to replicate tSCS findings (**Fig. 2, Extended Data Fig. 3)**, and demonstrated its utility for generating mechanistic hypotheses distinguishing cervical from lumbar tSCS (**Fig. 3, Extended Data Fig. 4,5).** The current model does not yet incorporate segment- and/or muscle-specific synaptic transmission or population-level phenotypic variability. We expect, however, that the proposed biophysical principles of volume conduction and axonal recruitment in cervical tSCS generalize across populations and muscles. The close alignment between simulated and experimental data supports the translatability of our virtual prototyping framework to other neurostimulation modalities engaging somatosensory afferents and motor efferents.

## Methods

### Computational framework

All simulations were performed with custom code developed in-house combining Sim4Life (version 7.2.1.11125, Zurich MedTech AG, Switzerland)^79^, Dolfinx (version 0.6, The FEniCS Project, UK)^80^, Python (version 3.11, Python Software Foundation, USA), and NEURON (version 8.4, Yale University, USA)^81^. Due to the extensive computational demands of this study, simulations were performed and distributed across multiple environments, preserving the same software versions. The environments included two high-performance computing (HPC) systems for large simulations and two local setups for smaller simulations. The first HPC system is the JUSUF cluster at the Jülich Supercomputing Centre, equipped with nodes having 2xAMD EPYC 7742 composed of 2×64 cores and running at 2.25 GHz and 256 (16×16) GB DDR4 of RAM. The second HPC system is the Meggie parallel cluster by NHR@FAU, equipped with two Intel Xeon E5-2630v4 "Broadwell" chips (10 cores per chip) running at 2.2 GHz with 25 MB Shared Cache per chip and 64 GB of RAM. The local setups included a workstation equipped with 20 Intel Xeon Gold 5218R cores running at 2.1 GHz with 512 GB of RAM and a compute server equipped with 2xAMD EPYC 9754 128-Core Processor running at 2.25 GHz with 1024 GB of RAM.

### 3D model of the human body

We developed a 3D model of the entire human body based on a previously published whole-body model (Jeduk version 4.1)^59^. As is, the model contained 1189 tissues that comprised the entire human body of a 33-year-old male adult, but lacked anatomically realistic volumes of the gray matter, white matter, cerebrospinal fluid, dura mater, epidural space, and spinal rootlets within the vertebral canal (**Fig. 1a**). We included all these tissues except for the spinal rootlets by generating them from an average spinal cord template of 50 healthy subjects (i.e. the PAM50 template)^60^. We started with extracting the centerline of the whole-body model’s spinal cord by computing center points of contours covering the craniocaudal axis of its white matter. We extracted these contours using the ‘CreatePlanarSlice’ function of Sim4Life’s XCoreModeling module starting from the top of the spine to the bottom with a step size of 10 mm. We then extracted the tridimensional points along these contours by converting the contour entity to a polyline using the ‘ConvertToPolyLine’ function and applying the ‘GetVertices’ function to the resulting entity. We then calculated the average of all contour points to retrieve the center point of each contour and interpolated the spinal cord’s centerline using SciPy’s ‘interpolate.splprep’ and ‘interpolate.splev’ functions to have a resolution of at least 2 points per millimeter of the spinal cord’s length. We then generated all volumes of the PAM50 template using ‘nii2mesh’^82^ command line tool. We then curved the resulting 3D structures to follow the centerline of the whole-body model’s spinal cord using the ‘curve modifier’ function in ‘Blender’ (version 3.6, Blender Foundation, Netherlands). The curved structures were then imported in Sim4Life^79^ and used to replace their counterparts in the Jeduk whole-body model.

To include anatomically realistic spinal rootlets, we developed a custom-code which populates the volumes between the peripheral nerves exiting the vertebral canal and the root entry zones (REZs) with curved cylindrical structures. For this purpose, we first defined the exit point of all rootlets belonging to the same spinal segment *s* merging into a peripheral nerve *P̄*_*s*,*Per*._ and the entry point of a given rootlet *r* belonging to segment *s* at the REZ *P̄*_*s*,*r*,*REZ*_. As the Jeduk whole-body model already included peripheral nerves, we defined the exit points of all rootlets corresponding to a single spinal segment as the peripheral nerve point proximal to their corresponding vertebrae with an offset of 0.5 mm from the dura mater. To define the entry point of a given rootlet into the white matter, we first had to define the segmental lengths and REZs along the craniocaudal extent of our spinal cord model. To do so, we calculated the relative percentage of segmental lengths for all segments from previously published morphometric data^61^, and applied these percentages to our spinal cord length (473.3 mm) to derive the length of each segment. We then calculated segment-specific ratios relating REZ length to segment length from previously published morphometric data^63^. We then defined the number of rootlets per spinal segment by scaling previously reported number of rootlets^63^ by the REZ length per segment in our model. We calculated rootlet diameters by dividing reported root diameters^63^ over the calculated number of rootlets in our model. We then defined the entry points for each group of rootlets corresponding to one segment by uniformly distributing these points along their corresponding REZ within the spinal cord’s centerline, resulting in *P̄*_*s*,*r*,*REZ*_ for each rootlet *r* and across segments of the spinal cord *s* ∈ *S*, *S* = {*C*1 … *S*4}. We then forced points along an algorithmically defined trajectory which represents the centerline of a given rootlet *r*. We defined this centerline trajectory of posterior rootlets to maintain a previously reported posterior/dorsal REZ entry angle (right: 40.1°, left: 39.8°)^62^, and the centerline trajectory of anterior rootlets to have an entry angle aligning with the direction of the ventral horn towards the center of the spine. To define the rootlet entry point into the white matter *P̄*_*s*,*r*,*REZ*(*WM*)_, we calculated the angle between each white matter contour point and the centerline at the craniocaudal level of *P̄*_*s*,*r*,*REZ*_ using numpy’s trigonometric functions. We then designated *P̄*_*s*,*r*,*REZ*(*WM*)_ as the point for which the calculated angle best matched the reported posterior/dorsal REZ angle^62^ and the angle aligning with the ventral horn for posterior and anterior rootlets respectively, using numpy’s ‘argmin’ function.

To define the trajectory of any given rootlet *r* belonging to segment *s*, we calculated an initial tridimensional set of points {*P̄*_*s*,*r*,*i*_}_*i*∈*I*_, *I* = {0 … *n* − 1}, using the REZ point *P̄*_*s*,*r*,*REZ*(*WM*)_ and the peripheral nerve exit point *P̄*_*s*,*Per*._ with equation (1):

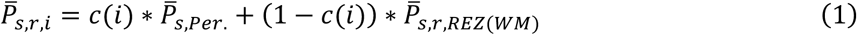

Here, *c*(*i*) = 1 − ^*i*^/_*n*_, whereby *i* is the index of a given point along the rootlet and *n* is the total number of desired points per rootlet, which we defined to be 3 points per millimeter. We then performed a for-loop using equation (1) from *i* = 0 to *i* = *n* to cover the trajectory from *P̄*_*s*,*Per*._ to *P̄*_*s*,*r*,*REZ*(*WM*)_. The resulting spline points were then processed to not collide with the white matter. To do so, we used ‘argmin’ function of numpy’s module to identify *P̄*_*s*,*r*,*i*,*contour*(*WM*)_ as the closest white matter contour point to *P̄*_*s*,*r*,*i*_ along its craniocaudal coordinate and *P̄*_*s*,*r*,*i*,*centerline*_ as the spine’s centerline point along the same craniocaudal coordinate. With *d*(*P̄*_1_, *P̄*_2_) being the distance between any two points (*P̄*_1_ and *P̄*_2_), we used ‘linalg.norm’ function of numpy’s module to calculate *d*(*P̄*^*s*,*r*,*i*^, *P̄*^*s*,*r*,*i*,*centerline*^) and *d*(*P̄*_*s*,*r*,*i*,*contour*(*WM*)_, *P̄*_*s*,*r*,*i*,*centerline*_). To quantify collision into the white matter, we then calculated *d*(*P̄*_*s*,*r*,*i*_, *P̄*_*s*,*r*,*i*,*contour*(*WM*)_) and multiplied it by -1 for all *P̄*_*s*,*r*,*i*_ having *d*(*P̄*_*s*,*r*,*i*_, *P̄*_*s*,*r*,*i*,*centerline*_) < *d*(*P̄*_*s*,*r*,*i*,*contour*(*WM*)_, *P̄*_*s*,*r*,*i*,*centerline*_). Accordingly, *d*(*P̄*_*s*,*r*,*i*_, *P̄*_*s*,*r*,*i*,*contour*(*WM*)_) was negative for all *P̄*_*s*,*r*,*i*_ colliding with the white matter. We additionally considered the collision resulting from the volume of the rootlet having a diameter (*RD*_*r*_) and thus defined the distance by which *P̄*_*s*,*r*,*i*_ must be pushed away from the spine’s centerline (*d*_*s*,*r*,*i*,*no col*._) with equation (2). We calculated *d*_*s*,*r*,*i*,*no col*_. only for *P̄*_*s*,*r*,*i*_ where 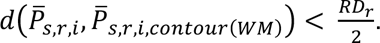

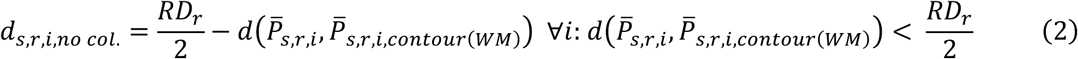

We then pushed all points {*P̄*_*s*,*r*,*i*_}_*i*∈*I*_ by the maximum *d*_*s*,*r*,*i*,*no* *col*._ across trajectory points (*d*_*s*,*r*,*max*.*no* *col*._) using equation (3)

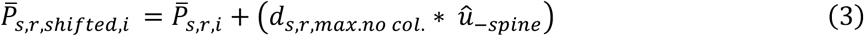

Here, *û*_–*spine*_ is the unit vector pointing away from the spine, which we calculated by subtracting *P̄*_*s*,*r*,*i*,*centerline*_ from *P̄*_*s*,*r*,*i*_ and dividing the resulting vector by its norm using numpy’s ‘linalg.norm’ function. We then ensured that the resulting trajectory {*P̄*_*s*,*r*,*shifted*,*i*_}_*i*∈*I*_ did not deviate from the originally defined peripheral nerve exit point by combining it with the original trajectory {*P̄*_*s*,*r*,*i*_}_*i*∈*I*_. To do this, we calculated a distance ratio *d*_*ratio*_(*i*) for each point as in equation (4) and used it to calculate the combined trajectory points {*P̄*_*s*,*r*,*combined*,*i*_}_*i*∈*I*_ using equation (5):

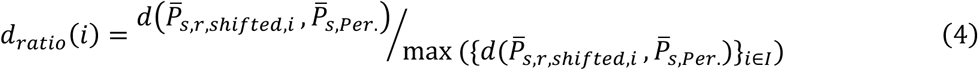

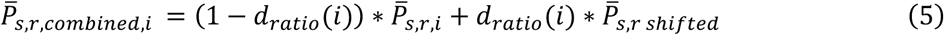

Applying equation (5) caused collision with the white matter for some trajectories. To avoid these collisions, we repeated the procedure described by equations (2) and (3) with {*P̄*_*s*,*r*,*combined*,*i*_}_*i*∈*I*_, resulting in {*P̄*_*s*,*r*,*refined*,*i*}*i*∈*I*_.

With *P̄*_*s*,*r*,*REZ*(*WM*)_ being the termination point in the trajectory {*P̄*_*s*,*r*,*refined*,*i*_}_*i*∈*I*_, we extended *P̄*_*s*,*r*,*REZ*(*WM*)_ towards the center of the spine by first calculating *P̄*_*s*,*r*,*REZ*(*GM*)_ as the entry point on the contour of the gray matter the same way described for *P̄*_*s*,*r*,*REZ*(*WM*)_. We then linearly sampled points from *P̄*_*s*,*r*,*REZ*(*WM*)_ to *P̄*_*s*,*r*,*REZ*(*GM*)_ using equation (6) to obtain the extension trajectory {*P̄*_*s*,*r*,*WM*→*GM*,j_}_j∈*J*_, *J* = {0 … *m* − 1}.

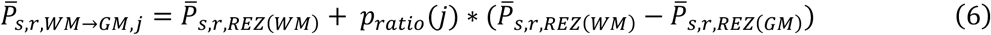

Here, *p*_*ratio*_(*j*) = ^*j*^/_*m*_, where *m* is the number of extension points that we computed to be the integer of the distance between *P̄*_*s*,*r*,*REZ*_(_*WM*_) and *P̄*_*s*,*r*,*REZ*_(_*GM*_) in millimeter. We then appended the points of {*P̄*_*s*,*r*,*WM*→*GM*,j_}_j∈*J*_ to {*P̄*_*s*,*r*,*refined*,*i*_}_*i*∈*I*_ to obtain the trajectory from *P̄*_*s*,*Per*._ to *P̄*_*s*,*r*,*REZ*_(_*GM*_) for each rootlet {*P̄*_*s*,*r*,*Per*.→*GM*,*k*_}_*k*∈*K*_, *K* = {0 … *n* + *m* − 1}. Rootlets of the sacral segments S3 and S4 entered the white matter with a larger craniocaudal distance between *P̄*_*s*,*r*,*REZ*(*WM*)_ and *P̄*_*s*,*r*,*REZ*(*GM*)_ compared to rootlets of other segments. To compensate for the large craniocaudal distance and to have a smooth extension for S3 and S4 segments, we extended *P̄*_*s*,*r*,*REZ*(*WM*)_ towards the spine’s centerline before using equation (3). To do this, we identified the closest spine’s centerline point along the same craniocaudal axis of *P̄*_*s*,*r*,*REZ*(*WM*)_ and added 20% of the direction vector towards the centerline to obtain the adjusted *P̄*_*s*,*r*,*REZ*(*WM*)_.

We aimed to extend rootlet trajectories into the gray matter. We thus extended *P̄*_*s*,*r*,*REZ*_(_*GM*_) in the same direction of *P̄*_*s*,*r*,*REZ*_(_*WM*_) to *P̄*_*s*,*r*,*REZ*(*GM*)_ by 20% the distance between *P̄*_*s*,*r*,*REZ*_(_*WM*_) and *P̄*_*s*,*r*,*REZ*_(_*GM*_), as this value produced trajectories just entering the gray matter without colliding with other trajectories across all segments. This resulted in the complete trajectory for each rootlet {*P̄*_*s*,*r*,*complete*,*l*_}_*l*∈*L*_, *L* = {0 … *q* − 1}, where *q* is the total number of resulting points along the trajectory. We avoided the extension beyond *P̄*_*s*,*r*,*REZ*_(_*GM*_) only for rootlets of sacral segments, since the gray matter had a relatively small cross-sectional area compared to cross-sectional areas of other segments, resulting in colliding trajectories. We then smoothed the resulting trajectory by interpolating points that double the trajectory’s resolution using SciPy’s ‘interpolate.splprep’ and ‘interpolate.splev’ functions.

We further introduced inter-root anastomoses between segments *s* ∈ *S* to replicate occurrences reported in a previous morphometric study of human cadavers^63^. For each anastomosis between any two segments *s*_*a*_ and *s*_*b*_, whereby *s*_*a*_ denotes the segment cranial to *s*_*b*_, we generated an anastomosis centerline. We designated each anastomosis centerline to start on the most cranial rootlet of *s*_*b*_ approximately 8 mm from *P̄*_*s*_*b*_,*r*,*REZ*_(_*GM*_) along {*P̄*_*s*_*b*_,*r*,*complete*,*l*_}_*l*∈*L*_ resulting in the anastomosis starting point (*P̄*_*s*_*b*_,*anstm*_) and to end 5 centerline points from *P̄*_*sa*,*r*,*REZ*_(_*GM*_) along {*P̄*_*sa*,*r*,*complete*,*l*_}_*l*∈*L*_ resulting in the anastomosis ending point (*P̄*_*sa*,*anstm*_). We then generated the centerline {*P̄*_*anstm*_(_*s*_*b*_→*sa*_)_,*u*_}_*u*∈*U*_, *U* = {0 … *v* − 1} using equation (7), whereby *v* is the number of points producing a resolution of 3 points per millimeter.

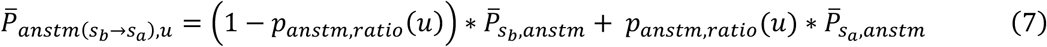

Here, *p*_*anstm*,*ratio*_(*u*) = (*u* + 1)/*v* represents the relative position along the anastomosis centerline. Using numpy’s ‘append’ function, we then concatenated {*P̄*_*s*_*b*_,*r*,*complete*,*l*_}_*l*∈*L*_, {*P̄*_*anstm*_(_*s*_*b*_→*sa*_)_,*u*_}_*u*∈*U*_ and {*P̄*_*sa*,*r*,*complete*,*l*_}_*l*∈*L*_ sequentially to form the complete centerline of the anastomosis. To allow for a smooth spline generation, we excluded some points from the trajectories before the concatenation. Specifically, we excluded *l* ∈ {*q* − 1}, *u* ∈ {1, *v* − 2, *v* − 1} and *l* ∉ {*q* − 2, *q* − 1} from {*P̄*_*s*_*b*_,*r*,*complete*,*l*_}_*l*∈*L*_, {*P̄*_*anstm*(*s*_*b*_→*s_a_*),*u*_}_*u*∈*U*_ and {*P̄*_*sa*,*r*,*complete*,*l*_}_*l*∈*L*_ respectively.

To build the complete rootlet centerlines trajectories extending to the peripheral muscles in the whole-body model, we extracted centerlines of the peripheral nerve trajectories from the whole-body model using ‘ConvertToPolyLine’ and ‘GetVertices’ functions of Sim4Life’s XCoreModeling module. We then concatenated rootlet centerlines to the respective peripheral nerve trajectories using numpy’s ‘append’ function. Upon concatenating the rootlet centerlines to the respective peripheral nerve trajectories, we noticed that some of the lumbosacral nerves exhibited steep bending angles below the pedicles. To avoid these steep bending angles, we removed 5 and 10 points from the rootlet and nerve centerlines respectively around the merging points where the two trajectories meet. We removed points only for segments where the craniocaudal distance between *P̄*_*s*,*r*,*REZ*(*GM*)_ and *P̄*_*s*,*Per*._ exceeded 20 mm and if both trajectories had enough number of points. We then used the ‘CreateSpline’ function of Sim4Life’s XCoreModeling module to generate the rootlet centerlines (**Fig. 1b**). Using the calculated rootlet diameters, based on previously reported measurements^63^, we then converted these rootlet centerlines into 3D rootlets using the function ‘ThickenWire’ of Sim4Life’s XCoreModeling module (**Fig. 1b**). To compensate for discontinuities observed in the original 3D structures of the peripheral nerves in the whole-body model, we supplemented them with cylindrical structures around peripheral nerve trajectories with 2.0 mm diameters, a value that guaranteed encapsulating all the trajectories.

### Electrode positioning

To realistically model skin-surface electrodes, we developed a python algorithm that automatically warps rectangular or circular surface electrodes with defined dimensions and a point of projection onto any 3D surface. To build rectangular electrodes, the algorithm takes as input a user-defined projection center point on the 3D model before slicing the projection surface with the ‘slice’ function of the Python library ‘PyVista’^83^ to retrieve the contours of the 3D model corresponding to a user-defined electrode dimension and shape. The algorithm then reconstructs the projected surface electrode from these retrieved contour lines using the ‘delaunay_2d’ function of PyVista^83^. For circular electrodes, the algorithm leverages the mesh interpolation function ‘interpolate’ from the Python library ‘PyVista’^83^ to reconstruct the electrode onto the projection surface by using the point of projection and the target radius. To ensure uniform resolution across all the different electrodes, we reconstructed electrode surfaces from grids of a uniform resolution (2 points/mm in each direction). We performed this reconstruction using the ‘RectilinearGrid’ class in ‘PyVista’^83^ to construct grids encapsulating the electrodes and ‘cKDTree’ in ‘SciPy’ to sample electrode points from the grids.

### Volume conduction

To use our 3D model in volume conduction simulations, we sought to discretize it into a structured mesh using the voxelization engine embedded within the ‘Electro Ohmic Quasi-Static’ simulation in Sim4Life^79^. To avoid having any holes or discontinuities in the generated voxels, we added a tissue filler volume and assigned it the lowest voxel priority setting and a coarse resolution. We generated this tissue filler by first voxelizing the skin as the volume encapsulating all tissues in the whole-body model with a fine grid (0.8 mm voxel size). Then we exported the voxels using the ‘WriteInputFile’ function in Sim4Life^79^, and used Pyvista’s^83^ ‘RectilinearGrid’ and ‘UnstructuredGrid’ classes, in addition to its ‘extract_surface’ function to create a surface mesh file (STL format) out of the skin voxels. We then imported the surface into Blender and used the ‘Boolean Modifier’ to subtract the skin surface from a bounding box encapsulating it, resulting in a filled inner volume representing the whole-body model. To make sure that this volume stays within the boundaries of the original skin, we used Blender to scale the resulting inner structure in the x and y directions with factors of 0.97 and 0.99 respectively, and exported it as an STL file to be imported again in Sim4Life^79^. We then specified tissue-specific voxelization parameters using the voxels settings of the ‘Electro Ohmic Quasi-Static’. We argued that including all tissues of the whole-body model in every simulation would massively increase computational demands without an added value in terms of accuracy. We thus defined paradigm-specific volume boundaries encapsulating only the electrode placements and all neural tissues being investigated within that paradigm by modifying the grid padding settings of the ‘Electro Ohmic Quasi-Static’ simulation in Sim4Life^79^. We then ran the voxelization engine to generate paradigm-specific meshes resulting in structured meshes composed of voxels in having a minimum of 95 and a maximum of 601 million voxels. To reduce computational demands and run simulations without the background voxels caused by the structured mesh generation in Sim4Life, we exported the discretized 3D model using the function ‘WriteInputFile’ in Sim4Life^79^. We then converted the resulting mesh into an unstructured mesh composed of hexahedrons using Pyvista’s^83^ ‘RectilinearGrid’ and ‘UnstructuredGrid’ classes, resulting in 64 to 448 million voxels that we then dumped into XDMF files using the ‘Mesh’ class of the Python module ‘meshio’^84^.

We assigned conductivity values (σ) to all anatomical structures based on an established publicly available material database^85^. We modified the skin conductivity in our model to replicate the electrode-skin interface of previously published computational tSCS models^35,36,68,69^, which was shown to satisfy bioimpedance measurements in a human tSCS study^35^. For the white matter, spinal rootlets, and peripheral nerves we assigned anisotropic conductivity values by leveraging a previously published methodology^5^, which leverages ‘Electro Ohmic Quasi-Static’ simulations in Sim4Life^79^ with Dirichlet boundary conditions assigned to the boundaries of nerve structures to retrieve the electric field tensors representing diffusion orientation inside the nerve structures. Conductivity tensors were then computed by aligning the conductivity of longitudinal and transversal components (σ_longitudinal_ = 7.33e-1 S/m and σ_transverse_ = 2.31e-1)^85^ with the resulting gradients from the simulations^5^.

We assumed perfect electrode conductivity and electrode-skin contact and used Dirichlet boundary conditions on the electrodes’ surfaces to simulate a potential difference between the two electrodes. To replicate electrodes’ polarity of stimulation devices used in the investigated tSCS paradigms^37,38^, Digitimer DS7A, and Digitimer DS8R (Digitimer Ltd, UK), we sought to assign values so that the cathode has a lower potential than the anode as stated in their respective user manuals. Therefore, we assigned V(x) = 0 and V(x) = 1 for the cathode and anode respectively. We considered zero-flux on the exterior boundaries to avoid hard cuts of the potential distribution on the boundaries. Similar to previously published computational models^5,24,35,36,68,69^, we solved the Laplace formulation of Maxwell’s equations shown in equation (8) by using an iterative solver in DolfinX^80^ resulting in electric potential obtained at each point of the discretized model.

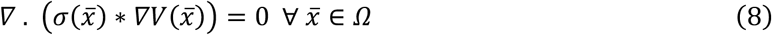

Here, *∇* . denotes the divergence, while *V*(*x̅*) is the electric potential and *σ*(*x̅*) is the conductivity tensor at any tridimensional coordinate *x̅* ∈ Ω, whereby Ω denotes the domain.

### Populations of axon trajectories

We populated the posterior roots, anterior roots, dorsal column, phrenic nerve, and vagus nerve, as well as all nerves of the brachial plexus and lumbosacral plexus with splines representing trajectories of axons. Posterior and anterior roots were populated with over 20,000, and 24,000 somatosensory afferent and motor efferent axons in the C5-T1 and L1-S4 spinal roots respectively. Based on a previously reported study showing a correlation between the number of axons and root diameters^86^, we populated the rootlets in our model with a number of axons that are proportional to their respective diameters. To have a minimum number of axons in any given rootlet, we first defined an arbitrary minimum number of axons (*min N*_*axons*_ = 3) to be populated in the rootlet with the minimum diameter (*min RD*) across all rootlets. For a given rootlet *r* with diameter (*RD*_*r*_), we then calculated the number of axons (*N*_*axons*,*r*_) using equation (9) to preserve the reported correlation^86^:

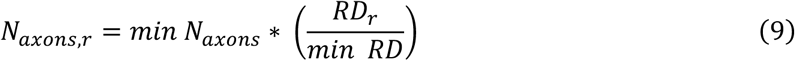

We then applied the same method to the phrenic nerve and the branches of the vagus nerve. To define the trajectories of axons, we leveraged the rootlet centerline trajectories ({*P̄*_*s*,*r*,*complete*,*l*_}_*l*∈*L*_), described in the subsection “3D model of the human body” and distributed axons around each centerline using a previously reported method of weighted superposition of points inside the rootlets’ cross-sections along the centerline’s trajectory^5^. The same method was applied to populate the branches of the vagus nerve with axon trajectories. For the phrenic nerve as well as nerves of the brachial plexus and lumbosacral plexus, we first used trajectory names and termination points to associate each peripheral nerve trajectory in the whole-body model to the spinal segment it belongs to. Using ‘ConvertToPolyLine’ and ‘GetVertices’ functions of Sim4Life’s XCoreModeling module, we created additional instances of each peripheral nerve trajectory corresponding to the number of rootlet trajectories of the assigned spinal segment. We then connected the pair of rootlet-periphery trajectories by using numpy’s ‘argmin’ function to find the closest points within each pair and cropping the trajectories with numpy’s slicing accordingly. We then concatenated the cropped trajectory points and generated a spline from the concatenated points using ‘CreateSpline’ function of Sim4Life’s XCoreModeling module. To populate the dorsal column with axon trajectories, we leveraged the ‘Electro Ohmic Quasi-Static’ simulation in Sim4Life^79^ similar to the simulations described in the subsection “Volume conduction” to obtain diffusion tensors inside the white matter. We then exported simulation results from Sim4Life^79^ and then used Pyvista’s^83^ ‘streamlines’ function on the resulting electric field distribution to compute streamlines following the curvature of the white matter. Using ‘random.choice’ function of numpy’s module, we then sampled 24 trajectories that were located inside the dorsal column to represent dorsal column axons.

### Conductance-based compartmental cable models and axon diameters

We aimed to simulate axonal responses of anatomically plausible populations of somatosensory afferents and motor efferents using NEURON. For this purpose, we translated all axonal trajectories into conductance-based compartmental cable models representing myelinated and unmyelinated axons.

For the myelinated axons, we assigned either the original MRG model^54^, the MRG-type model pair introduced by Gaines et al.^55^, or our newly-tuned MRG-type model. For each MRG model variant a set of diameter-dependent morphological properties must be defined to accurately estimate the axonal response of a given MRG-type model, as described by McIntyre et al. ^54^. To define these morphological properties, we used the interpolation functions introduced by Gaines et al.^55^. However, these functions did not extend to diameters < 5.7 µm. To rectify this shortcoming, we used previously published models of small-diameter myelinated axons^87,88^ in addition to ‘stats.linregress’ function from scipy’s python module to obtain linear interpolation functions for smaller diameters. We assigned myelinated axons to all axonal trajectories mentioned in the subsection “Populations of axon trajectories”.

Due to the lack of conclusive data^75–78^ on distinct diameter distributions for somatosensory afferents and motor efferents in different segments of the cervical spine, we used diameter distributions of the human median nerve^46^ for axons in the posterior roots and anterior roots of the cervical region, as well as the phrenic nerve branch of the C5 root (**Fig. 1d**). We used previously reported diameter distributions of the human L4 root to assign diameters of axons in the lumbosacral region^77^, representing axon groups of large somatosensory afferents and motor efferents. We used a distinct diameter distribution to assign diameters for axons in the vagus nerve branches^64^. We then reused the same vagus nerve diameter distribution to assign diameters of axons in the dorsal column, since reported values fall in the same range^90^ (**Fig. 1e**). Since the distribution used for posterior and anterior axons seemed to be bimodal^46^, we grouped axons by two groups, referred to as large-diameter group (8-20 µm) and small-diameter group (1-7.9 µm).

For the unmyelinated axons, we assigned the previously published ‘sundt’ model for each rootlet’s centerline extending to the peripheral nerves of the whole-body model within the cervical posterior roots^89^. Based on the diameter of the published unmyelinated axon model^89^, we assigned all unmyelinated axons a diameter of 0.8 µm^89^.

To discretize each trajectory into the different model-specific compartments, we first defined the number of compartments using the axonal spline length obtained by summing all consecutive points’ lengths using numpy’s ‘cumsum’ function and dividing it by the length of the single compartment. Using numpy’s ‘cumsum’ function, we then assigned cumulative length values for all points along the axonal trajectory and calculated the cumulative lengths of all compartments to be placed along the axon. Using a for-loop, we then looped over compartments to define their coordinates along the axon. For each compartment, we used numpy’s ‘argmin’ function to find the closest two points before and after the compartment using the calculated cumulative lengths. We then used numpy’s ‘linspace’ function to linearly interpolate points with a resolution of 0.01 µm between the identified two points encapsulating the compartment. We then calculated the cumulative lengths of the interpolated points to compare to the compartments’ cumulative lengths. Using numpy’s ‘argmin’ function, we then identified the start and end points that best matched the calculated cumulative length of the compartment. We then calculated the center point along each compartment by averaging the two points representing its boundaries. For unmyelinated axons, we used the same method to discretize trajectories into uniform compartments, each having a length of 50 µm, following the same segment length in the published model^89^. We then used a previously published method to initialize the different NEURON objects representing the axonal compartments and their ion channels^74^, that varied depending on the simulated axon model (**Fig. 1c**). For the ion channel parameters of the previously published models^54,55,89^, we refer the reader to the files available on the public model database (https://modeldb.science/3810, https://modeldb.science/243841 and https://modeldb.science/187473 for the MRG model^54^, the MRG-type model pair^55^ and the unmyelinated axon model^89^ respectively).

To simulate axonal responses to the applied stimulation, we used the ‘extracellular’ and ‘xtra’ mechanisms of NEURON to subject each compartment to the potential value along its coordinates resulting from the volume conduction simulation^74^. We obtained potential values at the center coordinates of each compartment using ‘RegularGridInterpolator’ available in the Python module ‘SciPy’. We then applied the stimulation pulse, scaled by a factor representing the stimulation amplitude (*V*), by linking it to the stimulus current of the ‘xtra’ mechanism and using ‘Vector.play’ in NEURON. We then ran NEURON simulations with an integration time step of 0.005 millisecond. Using ‘APCount’ and ‘record’ utilities of NEURON, we recorded the nodal action potentials and membrane potentials of the simulated axons. By analyzing the recorded results, obtained at each node of the axon, we identified whether, where, and when an axon was activated. We defined activation by detecting a propagating action potential (AP) at any node along the axon. We detected the propagating APs by finding a node *ni* resulting in *NAP*_*ni*_ = 1 and *T*_*ni*_ < *T*_*n*j_ ∀ *nj* ≠ *ni*, whereby *NAP*_*ni*_ and *T*_*ni*_ are the number of APs and the time of AP at node *ni* respectively and *nj* is any node along the axon that is not *ni*.

### Axon model tuning

We aimed to generate a pair of compartmental cable models that distinguish between somatosensory afferents and motor efferents. To this end, we implemented a tuning procedure designed to minimize the discrepancy between simulated and experimentally reported electrophysiological properties obtained during median nerve stimulation^44,45^. To reproduce the experimental conditions of multiple electrophysiological studies^44,45^, we first simulated median nerve stimulation using the arm of our whole-body volume conductor model **(Extended Data Fig. 1a).** We then mapped axon trajectories within the median nerve onto compartmental cable models. We based these compartmental cable models on the original MRG-model^54^, modified at the level of the fluted internode (FLUT) to include a fast potassium channel, following the recommendation of McIntyre et al.^54^. To enable tuning the models, we developed an error metric by combining multiple error metrics representing how well a given pair of models replicated key electrophysiological property differences between somatosensory afferents and motor efferents (**Extended Data Fig. 1b**). We first developed metrics that quantify the electrophysiological property differences between somatosensory afferents and motor efferents. More specifically, we addressed differences in strength-duration (SD) curves (equations 10 − 12), excitability recovery cycle (RC) (equations 13 − 15) and action potential (AP) widths (equation 16). For each metric, we calculated the relative error between the reported values^44,45,91^ and the values calculated from the simulated pair of models using equation (17). Finally, using w of each metric (equations 10 − 16), we calculated a combined error metric using a weighted summation of all metrics’ errors as shown in equation (18). RC ratios metrics had larger weights contributing to the combined error, since we observed a larger sensitivity of this metric compared to others upon tuning ion channel parameters.

Metrics:

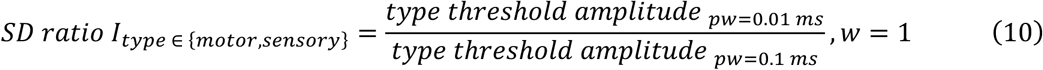

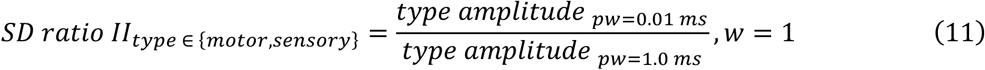

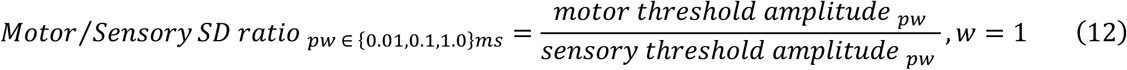

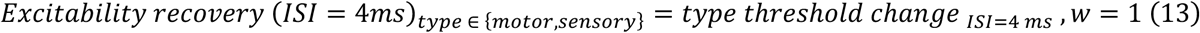

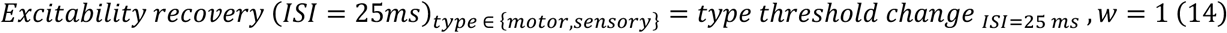

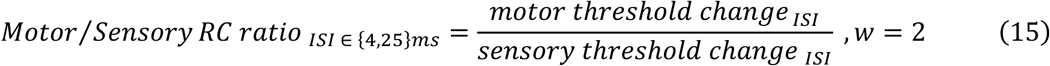

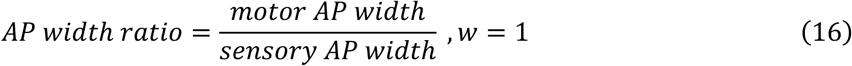

Errors:

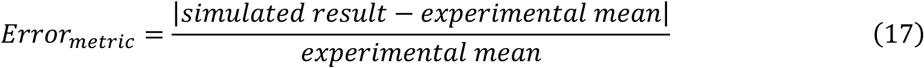

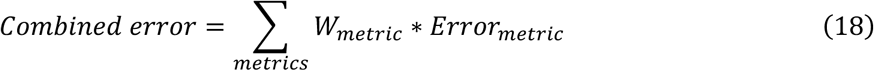

To construct pairs of somatosensory afferent and motor efferent models, we composed a parameter space spanning ranges of the ion channel parameters used in the MRG-type model. To enable efficient navigation of the large parameter space, we sampled sub-spaces to construct pairs of somatosensory afferent and motor efferent models. We implemented the subsampling by manually sampling specific values, falling in the range of predefined parameter ranges and creating permutations by nested for-loops over the chosen values across the different ion channel parameters. Using the potential distribution obtained from the volume conductor simulation of the median nerve stimulation (**Extended Data Fig. 1a**), we ran NEURON simulations to calculate the metrics and errors in equations (10 − 18), resulting in parameter combinations producing relatively low errors that we used as seeding points to the error minimization step (**Extended Data Fig. 1c**). We implemented error minimization using the Powell minimization method and basin-hopping as implemented in the Python module ‘SciPy’. Sub-space brute-forcing and error minimization resulted in multiple optima of pairs replicating the electrophysiological properties of somatosensory afferents and motor efferents (**Extended Data Fig. 1c**). We then chose one pair that resulted in an optimum strength-duration property fitting in the pulse width range relevant to this study (≥ 0.2 ms). We then compared our tuned MRG-type model pair with the performance of the previously published MRG model^54^ and the MRG-type model pair by Gaines et al.^55^ (**Extended Data Fig. 1d**).

### Recruitment curves

As we described in the subsection “Conductance-based compartmental cable models and axon diameters”, we ran simulations to calculate individual axons’ activation thresholds and to identify initiation sites of action potentials (ISAPs). For each simulated neurostimulation paradigm, activation thresholds were calculated by titrating the stimulus amplitude in repeated simulations until an action potential is initiated along the fiber with the smallest possible amplitude. We considered convergence when the relative change between activating and non-activating amplitudes is less than 1%. Axons were then grouped by their spinal roots or peripheral muscles to calculate recruitment curves of individual roots or individual muscles respectively, in addition to other groups such as the dorsal column, vagus nerve and phrenic nerve (**Fig. 3a-l, Extended Data Fig. 4a-l, 5a-d**). For a given neural structure *NrS*, we calculated the percentage recruitment value at an amplitude *V* using equation (19):

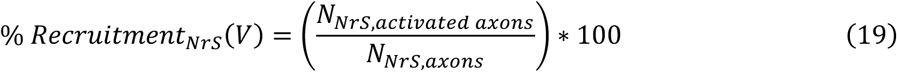

Whereby N_S,activated axons_ is the number of axons within the structure *NrS* that had an activation threshold ≤ V and N_*NrS*,axons_ is the total number of axons in that structure. In all the recruitment curves shown (**Fig. 3a-l**, **Extended Data Fig. 4a-l, 5a-d**), we included only axons with trajectories that could be traced and associated with individual upper limb muscles (**Fig. 3a-l**, **Extended Data Fig. 4a-l**) or lower limb muscles (**Extended Data Fig. 5a-d**). Across all NEURON simulations, we did not detect any activation of unmyelinated axons within the ranges of stimulation amplitudes. We therefore did not include the unmyelinated axons in the plotted recruitment curves.

### Motoneuron model

To simulate homonymous monosynaptic projections from somatosensory afferents onto motoneurons, we connected all myelinated motor efferent axons to motoneurons using a previously published motoneuron model^32^ (**Fig. 1f**). We generated 7265 and 14155 motoneurons connected to the (C5-T1) and (L1-S4) spinal root axons respectively. We adapted the morphological parameters of the motoneuron model by changing the soma dimensions to previously reported values (diameter = 53.04 µm)^92^ and by using a variable diameter of the initial segment that scales with the diameter of the axon, replicating a reported ratio of axon diameter to initial segment diameter (3.5/6.4)^93^. We used NEURON’s library in Python to initialize the motoneuron’s soma and initial segment for each efferent axon, following the methodology of a previously published code^32^. We then connected the initial segment of the motoneuron to the terminal node of the efferent axon trajectory {*P̄*_*s*,*r*,*complete*,*l*_}_*l*∈*L*_ upon entry into the gray matter (*P̄*_*s*,*r*,*REZ*(*GM*)_) using ‘Section.connect’ function of NEURON. Prior to each simulation, we added ‘APCount’ recorders in NEURON to the soma and initial segment of each motoneuron, enabling us to monitor APs of the motoneuron compartments, independent of the connected motor efferent.

### Synaptic transmission model and motor response classification

To simulate the assumed muscular response of the spinal cord to an artificially applied electric stimulus, we introduced a homonymous monosynaptic transmission model (**Fig. 1f** and **Extended Data Fig. 2**). To define the connectome of somatosensory afferent projections to motoneurons, we first created a mapping of somatosensory afferents and motor efferents to individual nerve branches and individual muscles. We based the mapping on nerve trajectory naming in the whole-body model and confirmed it by visually inspecting the innervation of the trajectories. We then initialized synaptic projections from large-diameter afferents^94,95^ to motoneurons belonging to the same peripheral nerve branch. In the case of multi-segment afferents belonging to the same nerve branch, we limited projections so that motoneurons receive inputs from afferents only in their entry segment and the adjacent segments just above and below their entry^65^. Each afferent input received on the motoneuron caused its membrane potential to increase by an excitatory postsynaptic potential (EPSP) which in turn raised the probability of initiating an AP depending on its value (**Extended Data Fig. 2b)**. Additionally, within the population of motoneurons, the percentage receiving afferent inputs (projection frequency) controlled the population-wise excitability by projections of somatosensory afferents (**Extended Data Fig. 2b)**. To simulate the afferent projections to motoneurons, we first simulated somatosensory afferents and then simulated motor efferents with the incoming afferent APs. We integrated afferent APs as excitatory synaptic inputs using their time of arrival at the respective motoneuron, which we achieved using the NEURON mechanism ‘vecevent’. For each afferent input, we created an excitatory synapse using the ‘ExpSyn’ point process in NEURON with a time constant = 0.5 ms^32^. Through the ‘NetCon’ object, we connected each event of spiking to the motoneuron’s soma using the ‘weight’ parameter. To assign a specific EPSP value to the motoneurons, we modified the ‘weight’ parameter to produce the chosen EPSP value for a diameter equal to the median across the simulated population. We then fixed the ‘weight’ parameter and assigned it to all motoneurons of that population. To assign a specific projection frequency, we uniformly sampled the percentage from motoneurons of the simulated population to be receiving somatosensory afferent inputs, while the rest had inactive synapses. We thus identified ranges of the EPSP and projection frequency from reported studies^49,50,52,53^ (**Extended Data Fig. 2b)** and tuned their capability in reproducing electrophysiological findings of upper and lower limb peripheral nerve stimulation studies^37,48^ **(Extended Data Fig. 2a)**. To do so, we replicated paradigms of peripheral nerve stimulation investigating the H-reflex of the flexor capri radialis (FCR) muscle^48^ and of the soleus muscle^37^ in our computational model **(Extended Data Fig. 2a)**. Upon running a simulation using a given EPSP value and projection frequency, we recorded AP times across nodes of each efferent axon as well as on the soma and initial segment of the respective motoneuron using ‘APCount.record’ in NEURON. We then used the AP times to classify APs along the efferents into afferent-initiated responses (AIRs) and efferent-initiated responses (EIRs). We started the classification by isolating different responses along an efferent axon using the AP times. We first considered the number of APs at the peripheral terminal of each efferent, as obtained from ‘APCount’ to be the number of responses of that efferent (*NAP*_*ni*=0_), whereby *ni* = 0 refers to the node at the peripheral terminal and *ni* = *LN* refers to the last node at the central terminal. For each response, we then processed APs of the consecutive nodes to determine which *AP*_*ni*c1_propagated into the current node’s *AP*_*ni*_ using AP time differences calculated with the ‘diff’ function in numpy’s module. We repeated this process for all responses along each efferent axon, resulting in an isolated temporal profile of APs per response. For each temporal profile, we then used the ‘argmin’ function of numpy to identify the node index associated with the earliest AP of that response (*ni*_*min*(*APTime*)_). If *ni*_*min*(*APTime*)_ was equal to the last node at the central terminal (*LN*) and was also preceded by an AP at the initial segment and soma of the motoneuron the response was considered to be resulting from the synaptic transmission and thus classified as an AIR. In all other cases the response was considered to be a result of directly activating the efferent axon and thus classified as an EIR. By summing up the number of AIRs (*NAIRs*) as well as the number of EIRs (*NEIRs*) of all efferents innervating a specific muscle across different stimulation amplitudes, we plotted recruitment curves exhibiting the characteristic shapes of H-reflex and M-wave curves^37,48^ (**Extended Data Fig. 2a**). From the calculated curves we calculated *AIR*_*max*_/*EIR*_*max*_ as the maximum *NAIRs* divided by the maximum *NEIRs* across all amplitudes (*V*), where *V* ranged from 0 to the amplitude able of activating all efferents innervating a given muscle (**Extended Data Fig. 2a**). We then compared *AIR*_*max*_/*EIR*_*max*_ to the reported *H*_*max*_/*M*_*max*_ ratios^37,48^ through calculating a percentage error (%*Error*_*H_max_* / *M_max_*_) using equation (20).

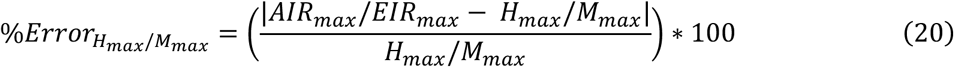

To tune the synaptic transmission parameters, we constructed a parameter space by sampling 25 combinations of the identified EPSP and projection frequency ranges. We then ran NEURON simulations with each of the parameter combinations and computed the percentage error using equation (20) (**Extended Data Fig. 2b**). Accordingly, we chose the parameter combination that resulted in a minimum %*Error*_*H_max_*/*M*_*max*__ of the FCR to be used for cervical monosynaptic projections and the combination that resulted in a minimum %*Error*_*H_max_*/*M*_*max*__ of the soleus to be used for lumbosacral monosynaptic projections across all simulated paradigms. For the tuning simulations, we used the paradigms that used a waveform of 1 ms in each of the reported studies^37,48^. We further evaluated the chosen monosynaptic transmission parameters in replicating findings of tibial nerve stimulation paradigms varying in stimulus waveforms, whereby we systematically varied the used axon models (**Extended Data Fig. 2c**).

### Motor threshold and the simulated percentage of afferent-initiated responses

Across simulated tSCS paradigms and for each simulated muscle, we defined AIR and EIR onsets as the minimum stimulation amplitudes (V) able of eliciting an AIR and EIR respectively. We then defined the minimum of the two onset amplitudes to be the motor threshold (MT) of the respective muscle. We used the minimum AIR and EIR onsets across all muscles for the values plotted in (**Fig. 3a-l**, **Extended Data Fig. 4a-l, 5a-d**). To quantify the contribution of somatosensory afferents to the elicited motor responses at any stimulation amplitude (V), we introduced the percentage of AIR (PAIR) metric that we calculated using equation (21). We leveraged PAIR by comparing its trends across different paradigms against trends of post-activation depression (PAD) reported in electrophysiological studies^37,38^ (**Fig. 2b-h, 6c,f,i,l, Extended Data Fig. 3e-h**)

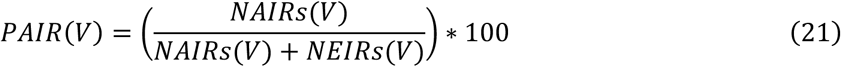

For all tSCS paradigms, we ran simulations with synaptic transmission across discrete stimulation amplitudes (V) ranging from 0 to a predefined value (*V*_*max*_) with a sampling step (*V*_*step*_). For tSCS paradigms that were used in comparisons against the reported PAD^37,38^ (**Fig. 2, Extended Data Fig. 3**), we defined *V*_*max*_ to correspond to the used experimental window with respect to MT^37,38^. For cervical tSCS paradigms shown in (**Fig. 5,6, Extended Data Fig. 8-10**), we initially defined *V*_*max*_ to be from 50 to 70 arbitrary units (a.u.) for different paradigms. We then modified *V*_*max*_ to cover MT of at least 6 out of 7 simulated upper limb muscles. In turn, for other cervical tSCS paradigms (**Fig. 3, Extended Data Fig. 4**) as well as lumbar tSCS paradigms (**Extended Data Fig. 5**), we only performed simulations until we detected both AIR and EIR onsets. We used a varying step (*V*_*step*_) ranging from 1 to 5 a.u. depending on the observed recruitments. Only when the difference between the AIR and EIR onsets was ≤ 2 a.u., we decreased the step of simulated amplitudes down to 0.25 a.u. in the range covering the onsets.

### Vagus nerve and phrenic nerve co-activation

We introduced the quantity vagus and phrenic nerve co-activation (VPNA) to quantify the unintended co-activation of peripheral structures unrelated to the facilitation of upper limb motor function (**Extended Data Fig. 8**). To calculate VPNA, we introduced equations (22) and (23), which we evaluated and averaged over amplitudes ranging from 0 to *min*(*V*_*max*_, 1.5 ∗ *MT*_*max*_), where *MT*_*max*_ was the maximum motor threshold across all muscles that elicited a response within the range [0, *V*_*max*_].

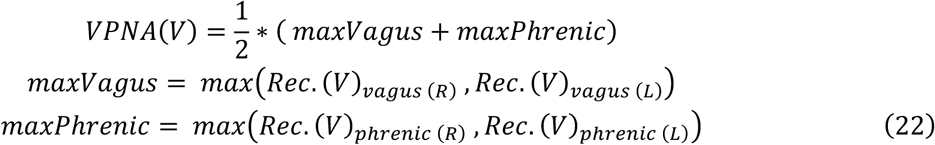

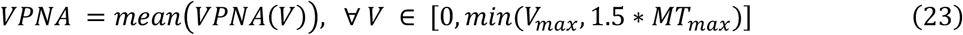

Here, *Rec*. (*V*)_*nerve*_ is the normalized recruitment of a given nerve at amplitude *V*. In equation (22), we denote right nerves by (R) and left nerves by (L).

### Electrophysiological Experiments

Our study combines previously published data on several participants (**Fig. 2, Extended Data Fig. 2,3**) with newly acquired electrophysiological experiments (**Fig. 6**). Please refer to the corresponding peer-reviewed publications for all relevant information regarding previously published data^37,38^, including statistical analysis and underlying assumptions. For all newly acquired data in our study, the experiments were reviewed and approved in July 2024 by the Ile de France III ethics committee (France Protocol Record: 2024-A00724-43, ClinicalTrials.gov Identifier: NCT07334977). 14 able-bodied participants (7 male, 7 female, average age 36.4 ± 13.3 years old) provided their informed consent to participate in this study. For each participant we tested three anode positions covering the distal to proximal end along the clavicles (**Fig. 6a**), C5C6 and C7T1 cathode position (**Fig. 6d**), with 32 mm circular, 50x50 mm square, and 50x70 mm rectangular anodes (**Fig. 6g**), and 0.2 ms, 0.5 ms, and 1.0 ms monophasic pulses (**Fig. 6j**). The tSCS stimuli were applied by delivering two monophasic pulses with an interstimulus interval of 50 ms of equal pulse amplitudes and using a constant current electrical stimulator (Digitimer DS7A, Digitimer Ltd, UK). The two evoked responses elicited by the stimuli were recorded at each muscle simultaneously. 16 Electromyogram (EMG) sensors were placed over the belly of upper arm and hand muscles bilaterally. The belly of each muscle was identified based on anatomical knowledge and confirmed by visually assessing at the EMG signal while the participant voluntarily contracted the muscle. Considered muscles were biceps brachii (BI), extensor carpi radialis (ECR), flexors carpi radialis (FCR), flexor digitorium superficialis (FDS), first dorsal interosseous (FDI), and adductor pollicis brevis (APB). EMG responses were recorded using a Bagnoli 16-channel system (Delsys Inc., USA) with a ground reference 50x70mm electrode placed over the left elbow of the participant. Before placing the electrodes, the skin was prepared using abrasive paste and then cleaned with alcohol. EMG signals were filtered between 20 and 450 Hz and amplified by 1000 using a 16-channel EMG amplifier (Bagnoli EMG system, Delsys Inc., USA). Analog outputs from the Bagnoli system were then digitized at 20 kHz through a data acquisition system (National Instruments NI 782258-0, USA, NI USB-6363, X-Series data acquisition module with BNC connectors). EMG data were acquired in a -50 to 150 ms window around each stimulus using a custom-built program in LabView (version 2024Q1, NI, USA). All data were subsequently processed offline in MATLAB (version R2024b, The MathWorks, Inc., USA). Stimulus pulses were delivered using an in-house developed Labview program. Stimulus amplitudes started from 5mA and were monotonically increased by steps of 5mA until the maximum tolerable amplitude for each participant. Motor threshold for an individual muscle was defined as the amplitude that evoked a visible EMG response to the first pulse of the paired stimuli in 2 out of 3 administered tSCS pulses.

In the previous tSCS electrophysiological studies used here^37,38^ as well as this study, PAD at a specific amplitude was calculated as the amount of depression in the second response (R2) compared to the first response (R1), PAD = (R2–R1)/R1. To have a fair comparison of our simulated PAIR to the measured PAD in previous tSCS electrophysiological studies^37,38^ (**Fig. 2b-h**, **Extended Data Fig. 3e-h**), we used the same amplitude ranges relative to MTs of respective muscles. In one study^37^, the range of 1x to 1.5x MT was used for each muscle and each paradigm when calculating PAD and the average over these amplitudes was reported. Therefore, we used the same ranges for calculating PAIR shown in **Fig. 2b-h**. In turn, for the other study^38^ the reported PAD values were averaged over amplitudes in the range of 1x to 2x the amplitude producing a visible PAD in a reference configuration (**Extended Data Fig. 3a**)^38^. We therefore modified the calculation of PAIR in those paradigms to be the average over a comparable range with the same reference configuration. Finally for PAD and PAIR of the newly conducted experiments, only values at MT of each muscle were considered (**Fig. 6c,f,i,l**). Statistical comparisons between tested tSCS paradigms with the newly acquired data were performed with a one-way ANOVA followed by a Bonferroni post-hoc correction using the Python library ‘statsmodels’. Our statistical analysis did not include additional covariates other than the parameters shown in **Fig. 6**.

## Acknowledgements

This work was partially supported by the “Bundesministerium für Forschung, Technologie und Raumfahrt” (German Federal Ministry of Research, Technology and Space - BMFTR) (project nr. 01ZZ2016 A.A., V.G., Z.H., and A.R.), the “Fondation pour la Recherche Médicale” (postdoctoral fellowship N.B.), the “Institut pour la Recherche sur la Moelle et l’Encephale” (IRME) (F.W. and N.B.), the “GPR inhibition program” (F.W.), the National Institutes of Health NINDS (award nr. K01NS127936 R.K. and I.S.), and the “Personalized Health & Related Technologies” program (PHRT 2022-279 E.N.). The authors gratefully acknowledge the FENIX Research Infrastructure for providing computing time on the FZJ Supercomputer JUSUF at Jülich Supercomputing Centre (JSC). The authors also gratefully acknowledge the scientific support and HPC resources provided by the Erlangen National High Performance Computing Center (NHR@FAU) of the Friedrich-Alexander-Universität Erlangen-Nürnberg (FAU). The hardware is funded by the German Research Foundation (DFG).

## Author contributions

A.A., U.H., K.M., F.W., A.R. conceptualized the tSCS mechanisms.

A.R. ensured funding, conceived and supervised all computational aspects of the study.

A.A., A.R. developed the computational framework.

A.A., E.N., A.R. developed the computational model.

A.A., V.G., Z.H. performed and analyzed computational simulations.

S.R.K, S.D. ensured HPC-compliance.

R.d.F., A.S., M.M., R.K., I.S. contributed electrophysiological datasets reported in two previously published tSCS studies, which supported model validation.

N.B., H.C., F.W. ensured funding, conceived, conducted, and analyzed data from the clinical study on tSCS mechanisms in 14 able-bodied individuals.

A.A., A.R. prepared the figures.

A.R. wrote the paper, and all the authors contributed to its editing.

## Data availability

Data from this study will be made available upon reasonable request to the corresponding author.

## Code availability

All software used to produce the figures in this manuscript will be made available upon reasonable request to the corresponding author.

## Extended data

**Extended Data Fig. 1.**
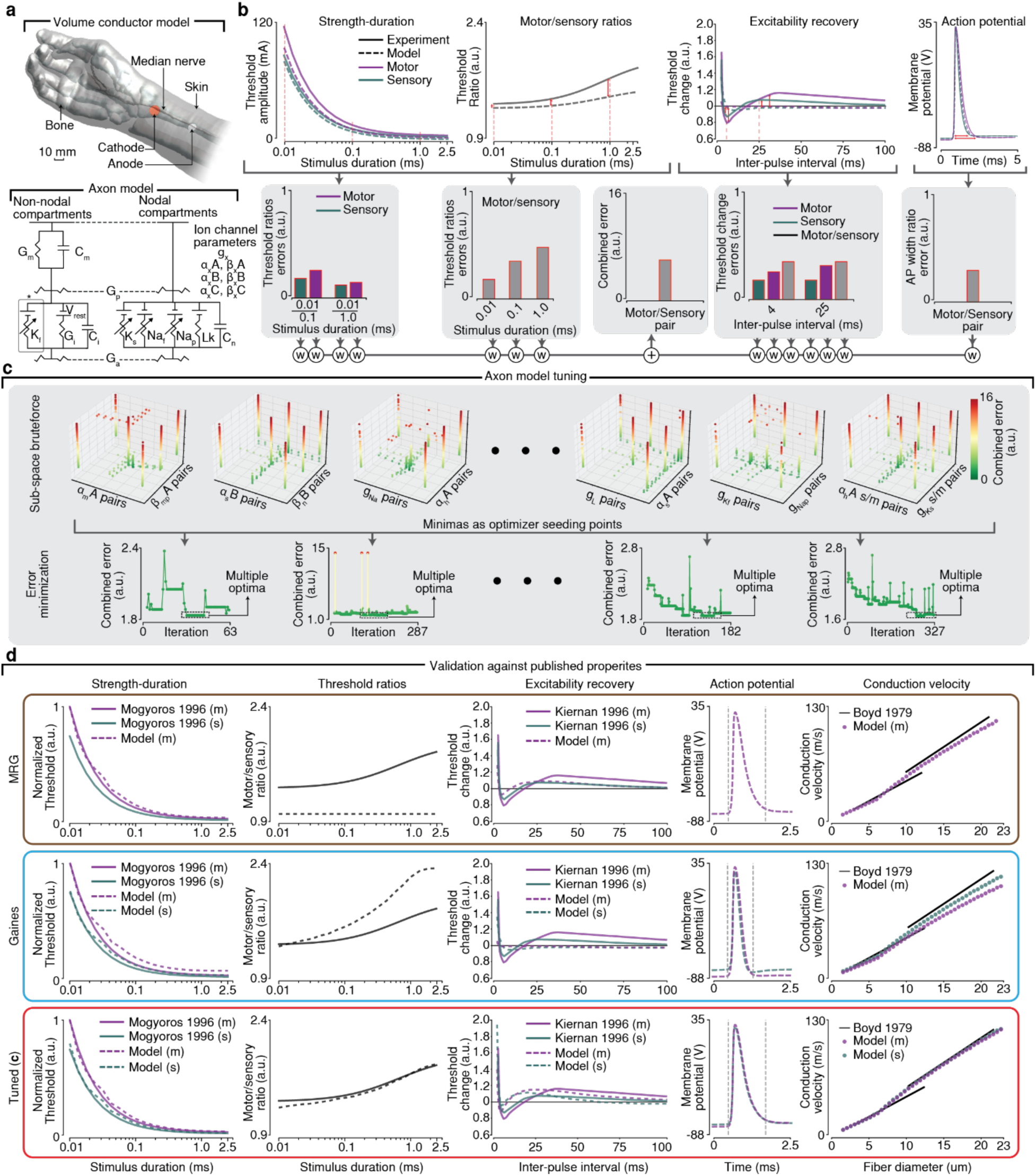
Process of tuning and validating a new MRG-type model pair. **a.** Replication of electrode placements in median nerve stimulation experiments^44,45^ in our computational model (top) to tune the parameters of the MRG-type model pair (bottom). *The fast potassium channel was only added to the fluted (FLUT) internodal segments. **b.** Comparison of model performance in replicating published electrophysiological properties of somatosensory afferents and motor efferents. Top: electrophysiological properties plotted against an example pair of models (left to right: strength-duration curves^44^, threshold ratios^44^, excitability recovery^45^ and action potential shape^91^). Bottom: Error metrics computed from each of the electrophysiological properties are weighed to compute a combined error used for optimization. **c.** We tuned the MRG-type model pair by combining two methods. Top: We performed sub-space brute-forcing by choosing combinations of somatosensory/motor values of the different model parameters that satisfy experimentally-reported differences^44,45,91^ and calculated a combined error for each parameter set representing how closely the models replicate the electrophysiological properties in (**b**). Bottom: Pair sets resulting in low errors from the brute-forcing steps were chosen as starting points for a minimization algorithm. The tuning resulted in multiple optimum sets of parameters reducing the combined error. From these we chose one that resulted in top strength-duration property fitting in the pulse width range relevant to this study. **d.** Comparison of the performance of previously published models in replicating the properties in (**b**). Top: MRG-type motor model^54^. Only the motor variant is shown since the MRG-model did not include a sensory variant and therefore motor/sensory ratio is set to 1 referring to modeling the sensory variant to be the same as the motor MRG. Middle: Gaines MRG-type model pair^55^. Bottom: Our tuned MRG-type model pair resulting from (**c**). Abbreviations: millimeter (mm), milliampere (mA), arbitrary units (a.u.), millisecond (ms), Volt (V), myelin conductance (G_m_), myelin capacitance (C_m_), periaxonal conductance (G_p_), axoplasmic conductance (G_a_), slow potassium (K_s_), fast potassium (K_f_), leakage (Lk), internodal conductance (G_i_), internodal capacitance (C_i_), resting membrane potential (V_rest_), persistent sodium (Na_p_), fast sodium (Na_f_), nodal capacitance (C_n_), ion channel conductance (g), ion channel opening rate (α), ion channel closing rate (β), ion channel maximum speed of transition (A), ion channel half-activation voltage (B), ion channel response slope (C), weight (w), sensory (s), motor (m).

**Extended Data Fig. 2.**
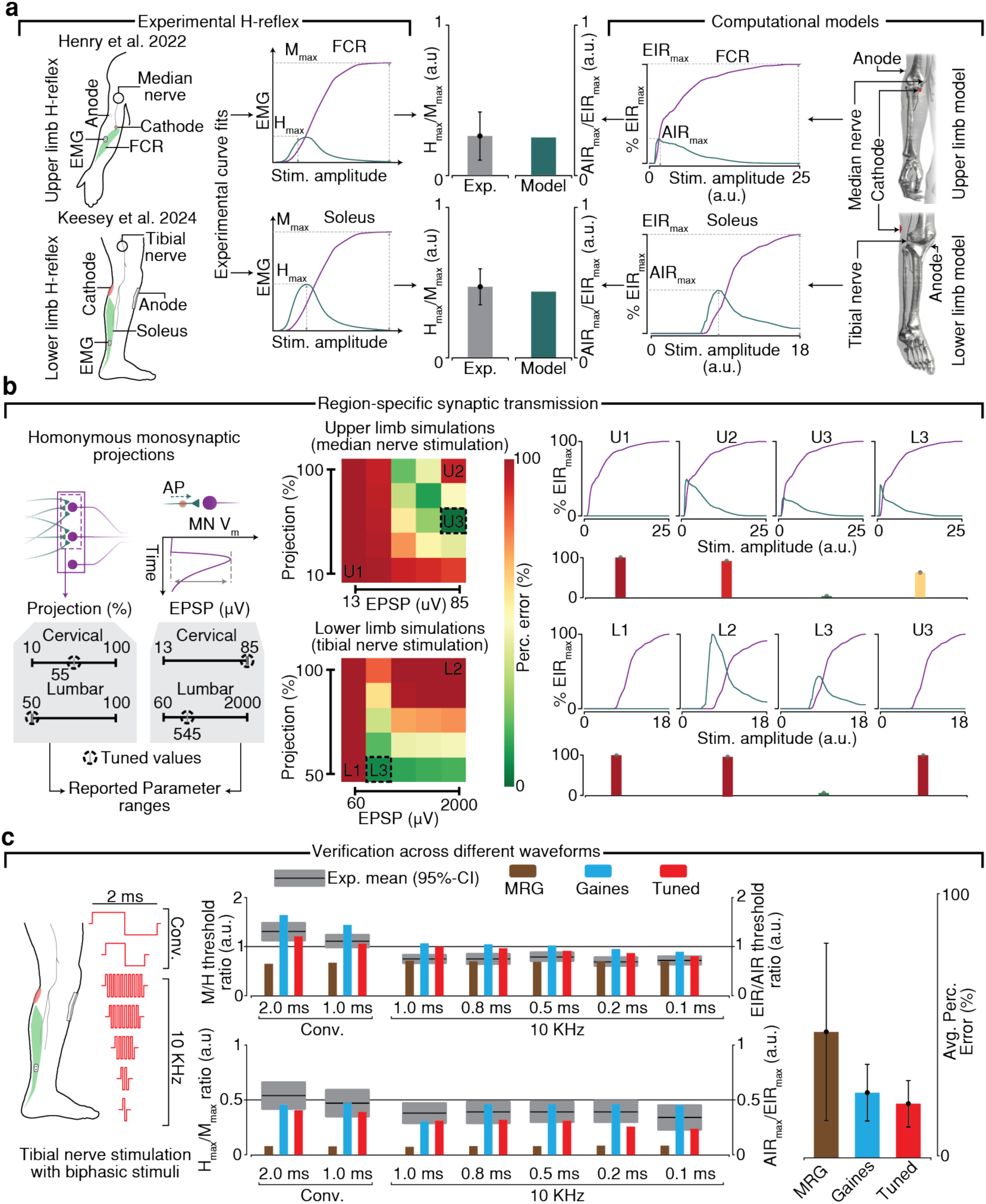
Process of tuning and verification of the homonymous monosynaptic transmission model applied to cervical and lumbar spinal roots. **a.** Left: Reported findings of H-reflex and M-wave recruitment curves in an upper limb muscle^48^ (top) and a lower limb muscle^37^ (bottom) were used as objectives for the tuning process, leveraging the reported H_max_/M_max_ ratios. Right: Replication of the experimental setups in each of the studies shown on the left in our computational model and the resulting efferent-initiated responses (EIR) and afferent-initiated responses (AIR) recruitment curves and AIR_max_/EIR_max_ ratio. **b**. Left: Reported values in electrophysiological studies of homonymous monosynaptic projections from Ia afferents to cervical motoneurons^49,53^ and lumbar motoneurons^50,52^ were leveraged to define the parameter space of our region-specific synaptic transmission model. Middle: Percentage error between the reported H_max_/M_max_ and the simulated AIR_max_/EIR_max_ ratios across discrete points of the parameter space for cervical (top) and lumbar (bottom) roots with dashed boxes highlighting the optimum parameters producing the lowest errors. Right: AIR and EIR recruitment curves of different points in the parameter space and their respective error values for cervical (top) and lumbar (bottom) roots. **c.** Verification of the resulting synaptic transmission model across different waveforms used in a tibial nerve stimulation experiment^37^ (Left). The reported M/H threshold and H_max_/M_max_ ratios were used to compare against EIR/AIR threshold and AIR_max_/EIR_max_ ratios respectively using each of the MRG-type model pairs (middle). Our tuned MRG-type pair produced the lowest average percentage error across the reported metrics. Abbreviations: electromyography (EMG), flexor capri radialis (FCR), maximum (max), excitatory post-synaptic potential (EPSP), microvolt (μV), percentage (Perc.), experiment (Exp.), confidence interval (CI), stimulus (stim.), arbitrary units (a.u.), millisecond (ms), conventional (Conv.), kilohertz-frequency (KHF), average (Avg.)

**Extended Data Fig. 3.**
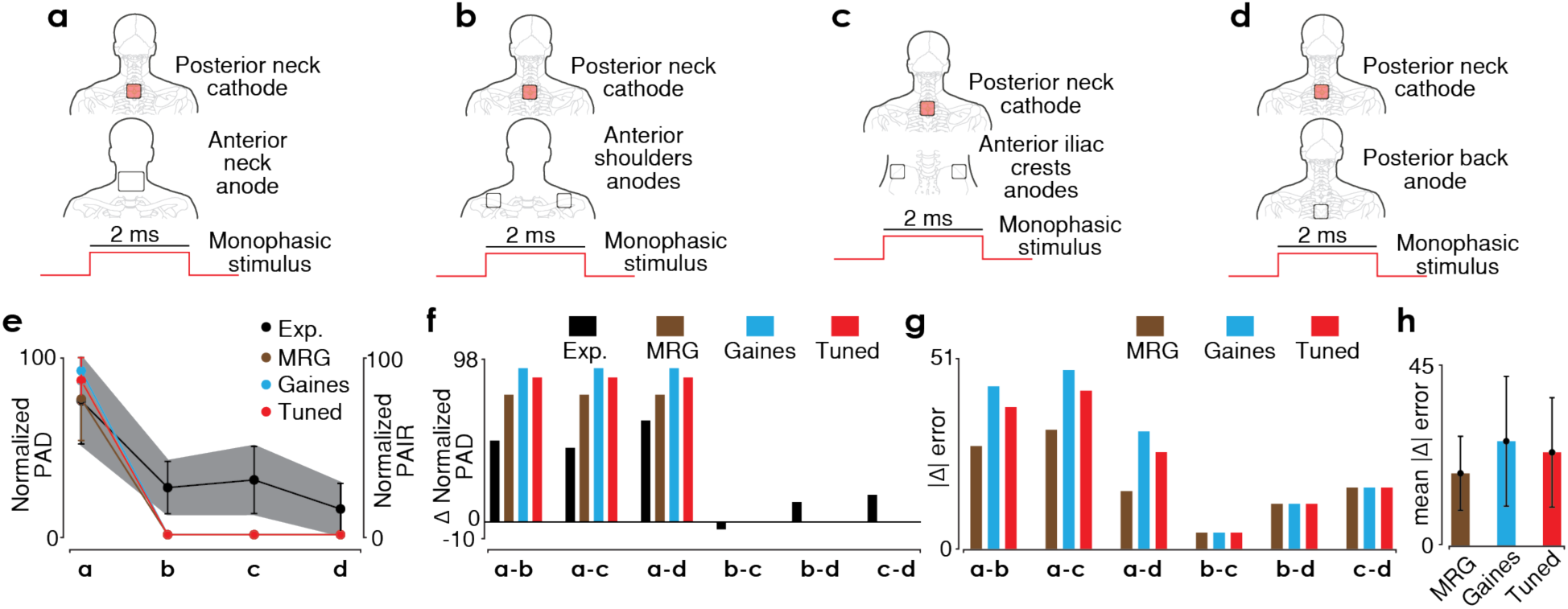
Validation of the computational model against PAD findings in a previously reported electrophysiological study of cervical tSCS^38^. **a-d.** Sketches of the applied tSCS paradigms in the study. **e**. Comparison of normalized post-activation depression (PAD) (black line represents the average value whereas the gray area represents the standard deviation) and Percentage of afferent initiated responses (PAIR) (colored lines) across the different tSCS paradigms from (**b-d**). **f**. Differences in PAD between tSCS paradigms (black bar) and differences in PAIR between tSCS paradigms (colored bars). **g**. Differences between PAD and PAIR across trends comparing tSCS paradigms. **h.** Average and standard deviation of **g** across trends of tSCS paradigms. Abbreviations: millisecond (ms).

**Extended Data Fig. 4.**
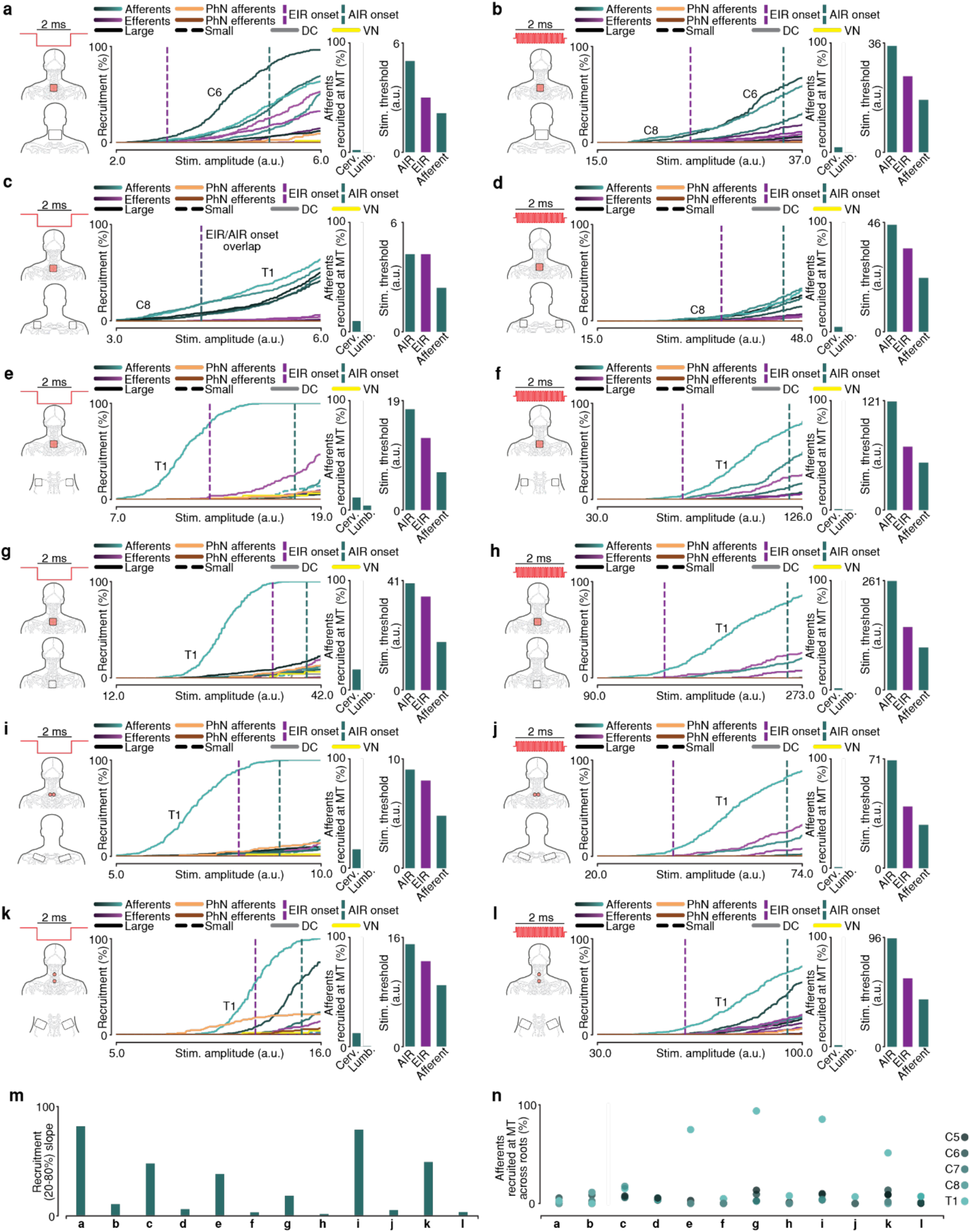
Even with inverted monophasic polarity or kilohertz-frequency (KHF) waveforms, the electrode placements from Fig. 3 yield lower activation thresholds for somatosensory afferents than motor efferents in silico, while motor thresholds remain dominated by efferent-initiated responses. **a-l** Stimulation configurations (left), recruitment curves for all simulated axons including afferent-initiated (AIR) and efferent-initiated response (EIR) thresholds (middle), and stimulation thresholds for AIRs, EIRs, and afferent recruitment (right) for six commonly used cathode and anode placements. Inverted monophasic stimulation is shown in (**a,c,e,g,i,k**), and KHF stimulation in (**b,d,f,h,j,l**). **m.** Recruitment slopes of somatosensory afferents across all cervical tSCS paradigms (**a–l**), are shown as an indicator of variance in the efficacy of somatosensory afferent recruitment relative to stimulation amplitude. **n**. Somatosensory afferents recruited at MT across posterior roots for all cervical tSCS paradigms (**a-l**) are depicted as an indicator of variance in the recruitment of somatosensory afferents between spinal roots. Abbreviations: phrenic nerve (PhN), dorsal column (DC), vagus nerve (VN), cervical (Cerv.), lumbar (Lumb.), motor threshold (MT), stimulus (stim.), arbitrary units (a.u.), millisecond (ms)

**Extended Data Fig. 5.**
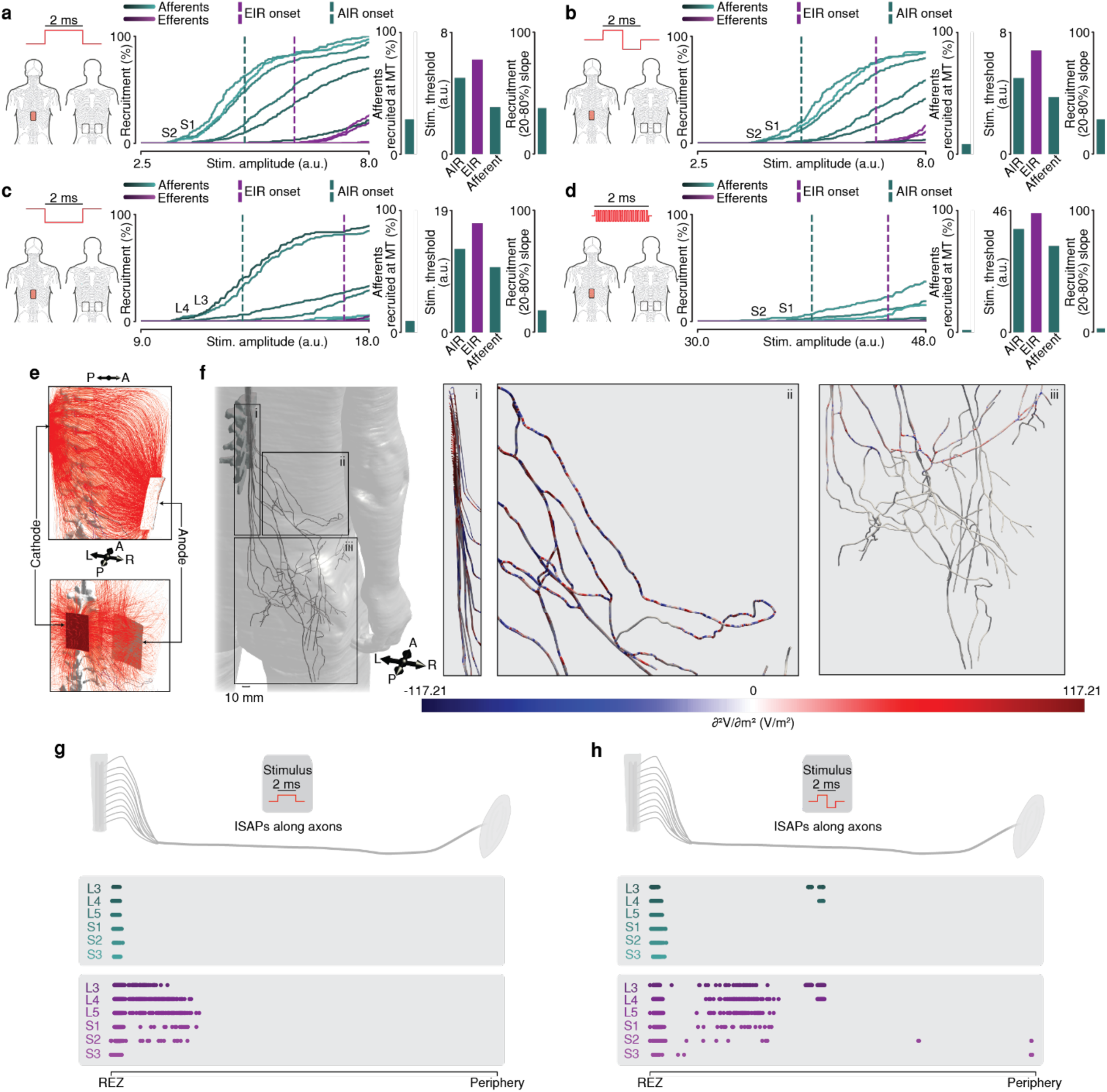
Both somatosensory afferents and afferent-initiated responses exhibit lower stimulation thresholds than motor efferents and efferent-initiated responses in lumbar tSCS^37^. **a-d.** Stimulation configurations (left), recruitment curves for all simulated axons including afferent-initiated (AIR) and efferent-initiated response (EIR) thresholds (middle), and stimulation thresholds for AIRs, EIRs, and afferent recruitment, as well as recruitment slopes (right) for lumbar tSCS with monophasic (**a**), biphasic (**b**), inverted monophasic (**c**), and kilohertz-frequency (**d**) stimulation. **e.** Electric field streamlines. **f.** Second spatial derivative of the extracellular potential along the axons (¶^2^V/¶X^2^) governing neural recruitment from the REZ to the periphery for lumbosacral roots using a monophasic stimulus. **g,h.** Initiation sites of action potential distributions for somatosensory afferents (turquoise) and motor efferents (purple). Abbreviations: motor threshold (MT), stimulus (stim.), arbitrary units (a.u.), millisecond (ms), anterior (A), posterior (P), right (R), left (L), millimeter (mm), initiation sites of action potentials (ISAPs), root entry zone (REZ), volt (V), meter (m).

**Extended Data Fig. 6.**
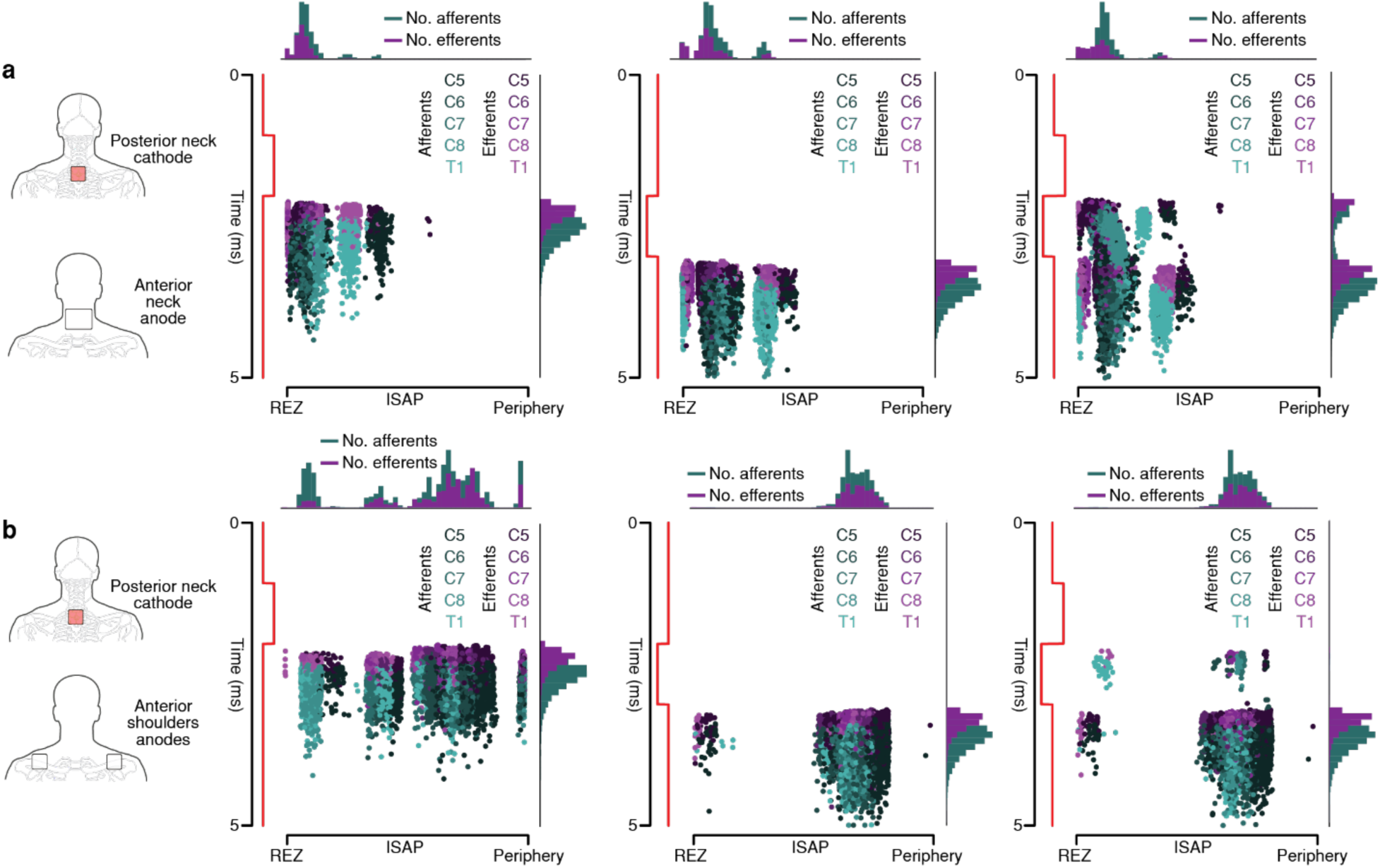
Negative phases of multiphasic waveforms induce action potentials along peripheral nerves for distal tSCS. **a.** tSCS with a posterior square (50x50 mm) cathode and an anterior rectangular (75x100 mm) anode on the neck and the resulting initiation sites of action potentials (ISAPs) with a monophasic positive stimulus (left), a monophasic negative stimulus (middle) and a biphasic stimulus (right). **b.** tSCS with a posterior square (50x50 mm) cathode and a pair of anterior square (50x50 mm) anodes on the shoulders and the resulting ISAPs with a monophasic positive stimulus (left), a monophasic negative stimulus (middle) and a biphasic stimulus (right). Abbreviations: millisecond (ms), number of (No.), root entry zone (REZ), initiation site of action potentials (ISAP).

**Extended Data Fig. 7.**
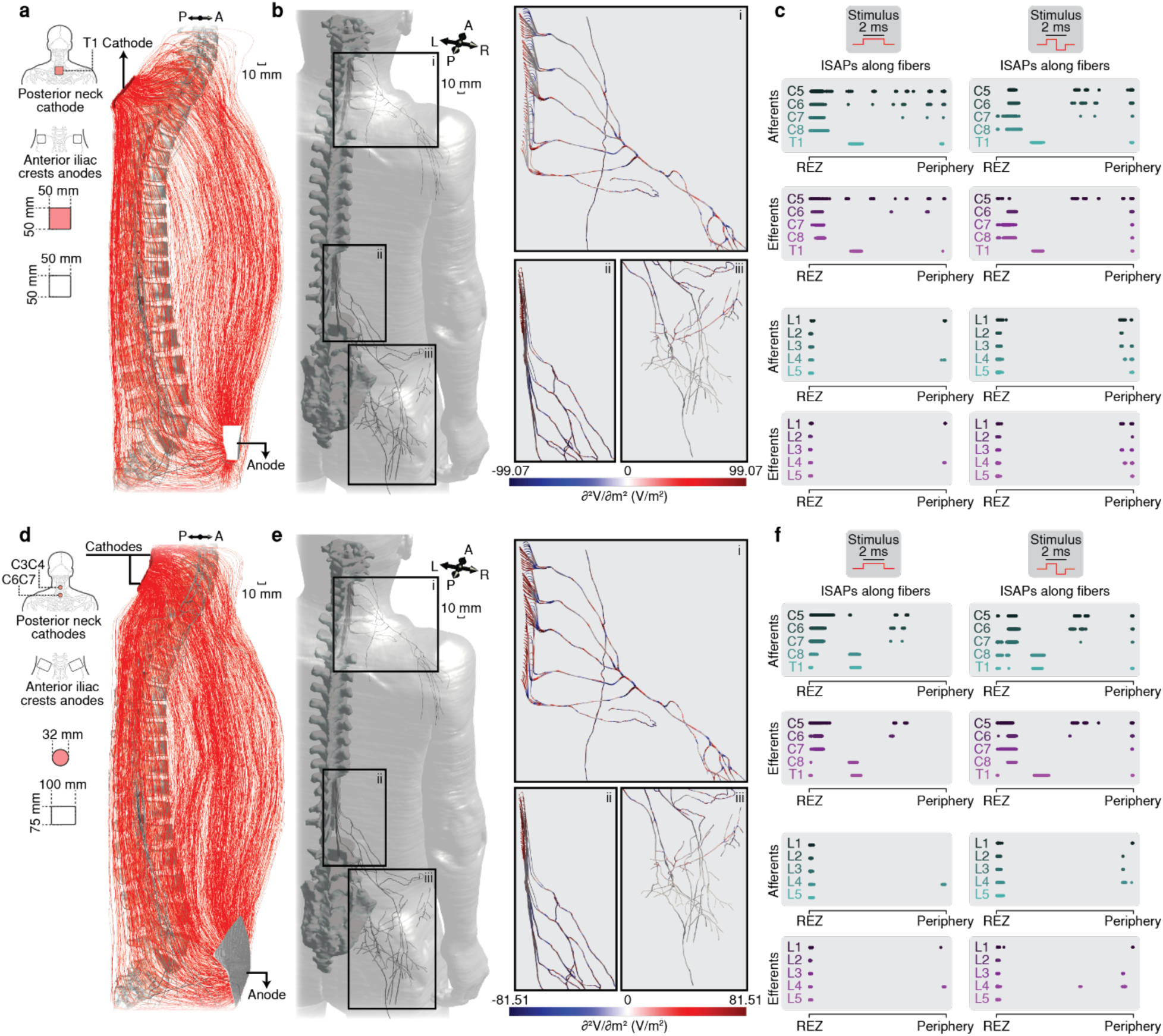
Cervical tSCS with iliac crest anodes can co-activate lumbar axons. **a.** Sketch showing the stimulation paradigm with the T1 square (50x50 mm) cathode and the two iliac crests square (50x50 mm) anodes (left) and the corresponding electric field streamlines shown with a sagittal view (right). **b.** Second spatial derivative of the extracellular potential along the axons (¶^2^V/¶X^2^) governing neural recruitment for cervical (i) and lumbar (ii and iii) roots corresponding to the stimulation paradigm shown in **(a)**. **c.** Initiation site of action potentials (ISAPs) for somatosensory afferents (turquoise) and motor efferents (purple) of cervical roots (top) and lumbar roots (bottom) using a monophasic stimulus (left) and a biphasic stimulus (right) corresponding to the stimulation paradigm shown in **(a)**. **d.** Sketch showing the stimulation paradigm with the two vertically aligned circular (32mm-diameter) cathodes and the two iliac crests rectangular (100x75 mm) anodes (left) and the corresponding electric field streamlines shown with a sagittal view (right). **e.** Second spatial derivative of the extracellular potential along the axons (¶^2^V/¶X^2^) governing neural recruitment for cervical (i) and lumbar (ii and iii) roots corresponding to the stimulation paradigm shown in **(d)**. **f.** Initiation site of action potentials (ISAPs) for somatosensory afferents (turquoise) and motor efferents (purple) of cervical roots (top) and lumbar roots (bottom) using a monophasic stimulus (left) and a biphasic stimulus (right) corresponding to the stimulation paradigm shown in **(d)**. Abbreviations: millimeter (mm), anterior (A), posterior (P), right (R), left (L), initiation sites of action potentials (ISAPs), root entry zone (REZ), volt (V), meter (m).

**Extended Data Fig. 8.**
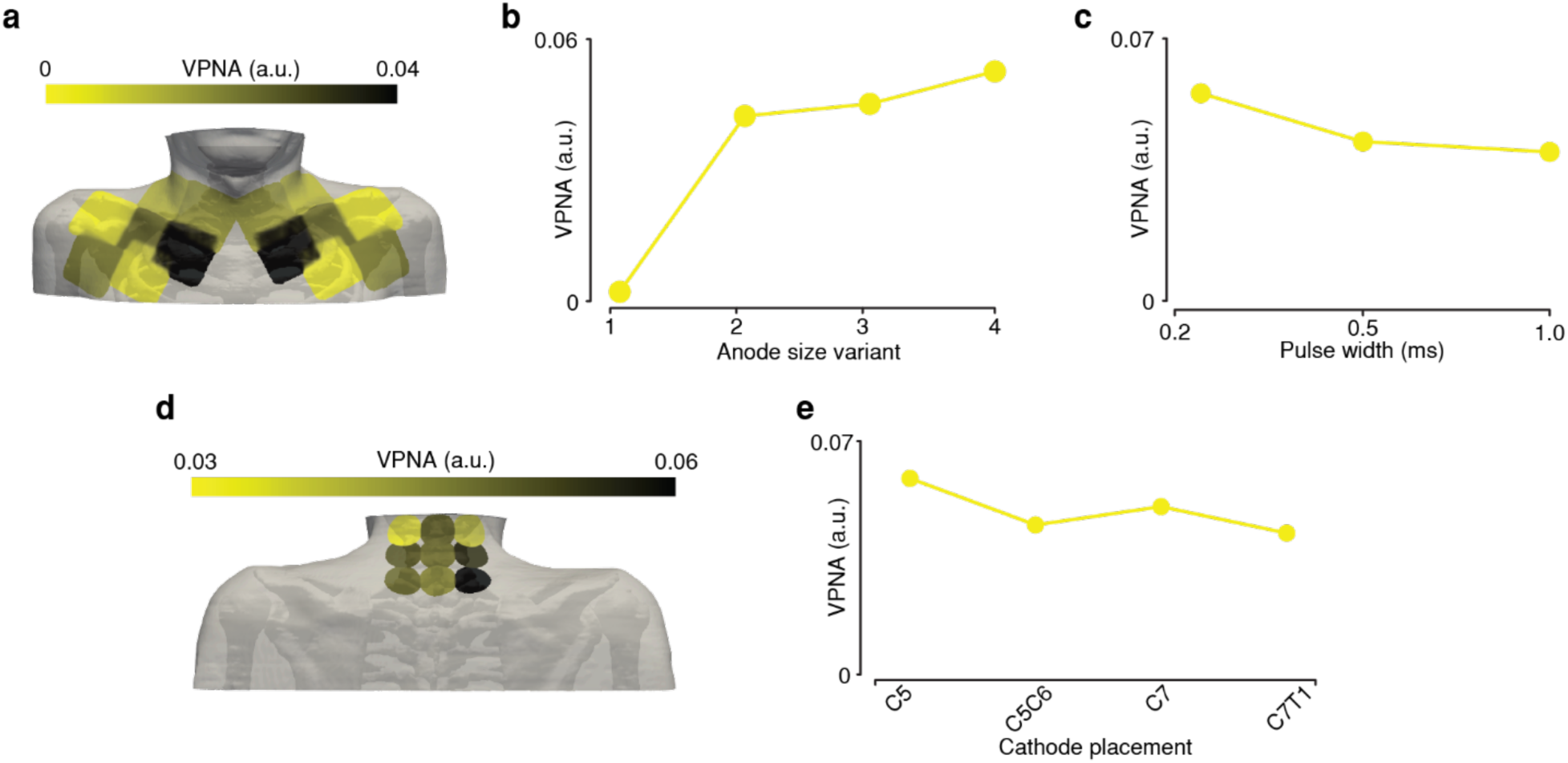
Minimal overall vagus and phrenic nerve co-activation (VPNA) across cervical tSCS paradigms and segment-specific behavior of cathodal placement. **a.** VPNA across the different anode placements shown in (Fig. 5a) for which an increase of co-activation for medial electrodes can be observed despite the overall small values. **b.** VPNA across the different anode sizes shown in (Fig. 5d) for which only the small circular anode (32mm-diameter) resulted in a VPNA of 0 with an increasing co-activation with larger anodes. **c**. VPNA for the different pulse widths shown in (Fig. 5g) with overall minimal values. **d**. VPNA across the different cathode placements shown in (Fig. 5j) showing a segment-specific and right-left variation of co-activation. **e**. VPNA for the different cathode placements in (Fig. 5m) with overall minimal values. Abbreviations: arbitrary units (a.u.), millisecond (ms).

**Extended Data Fig. 9.**
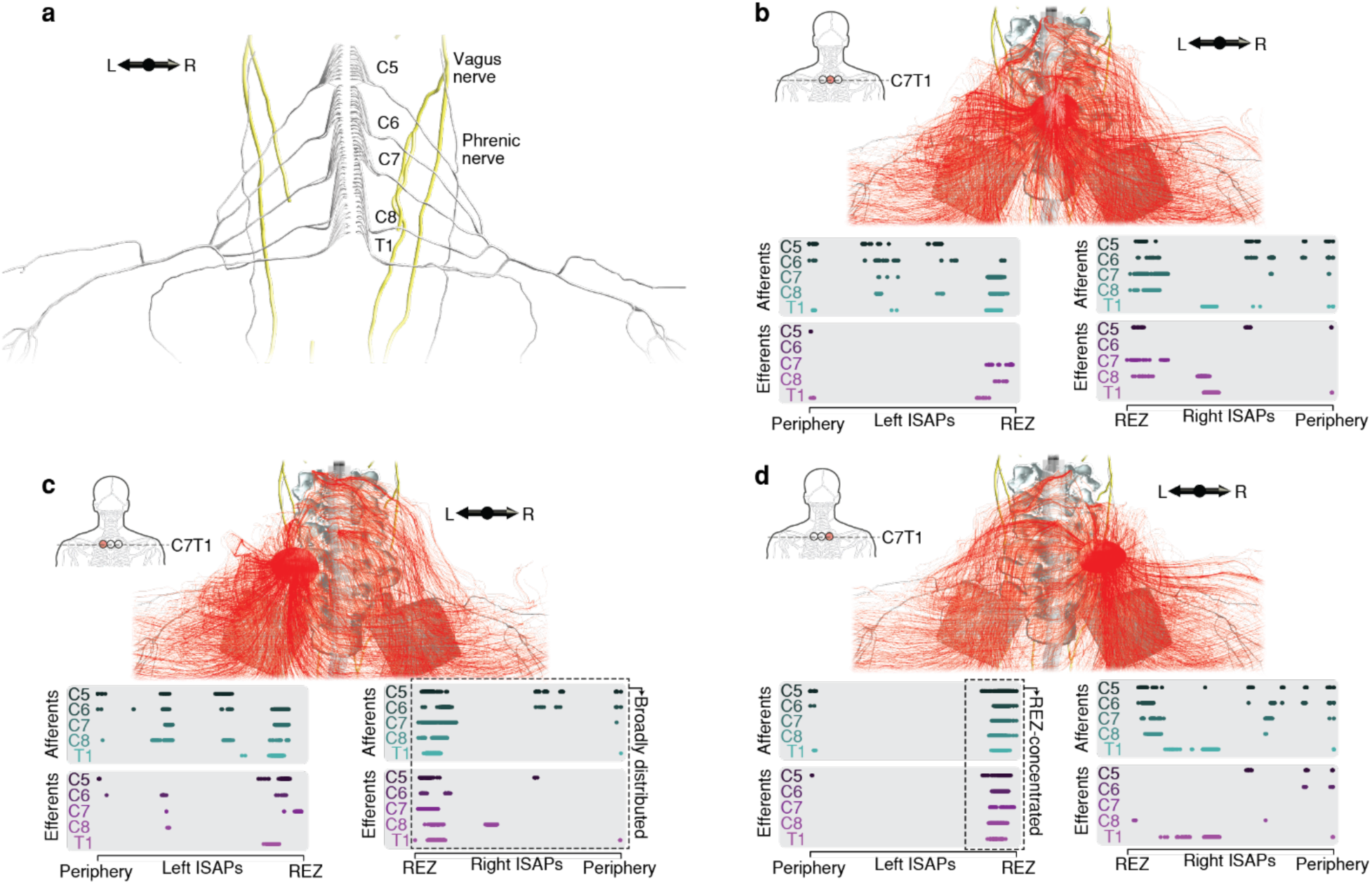
Asymmetries in nerve trajectories and stimulus currents cause different initiation sites of action potentials (ISAPs) for different cathode placements. **a.** 3D geometry of the right and left spinal roots and their peripheral nerve branches exhibit asymmetric trajectories in our model. **b-d.** Stimulus current trajectories resulting from three cathode placements sampled from the most caudal row of the grid shown in (Fig. 5j) and the resulting ISAPs for right and left roots. The right cathode in (**d**) caused ISAPs of the contralateral (left) roots to be concentrated around the REZ, while the left cathode in (**c**) did not have the same effect on right roots. Abbreviations: right (R), left (L), root entry zone (REZ).

**Extended Data Fig. 10.**
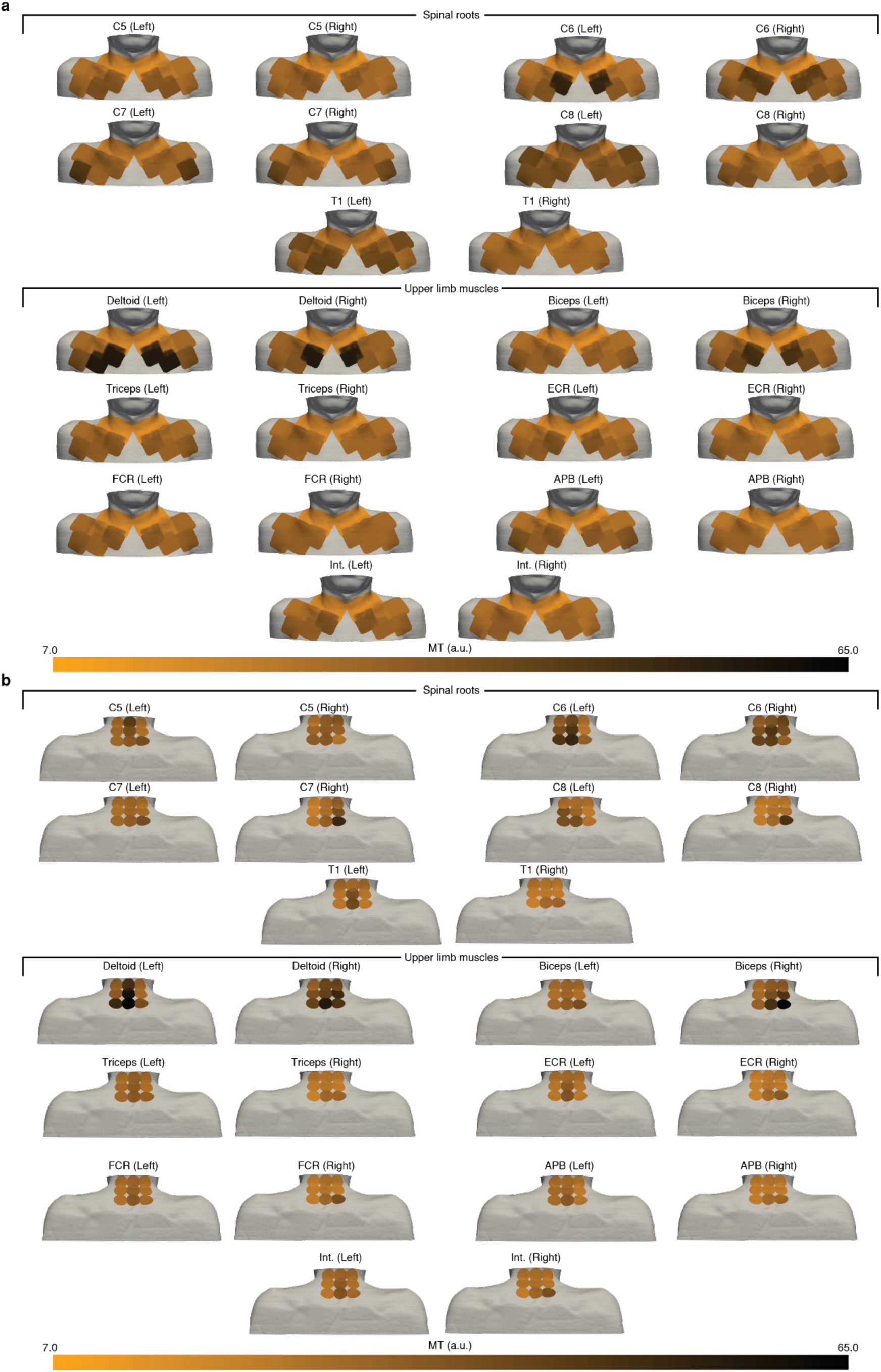
Motor threshold (MT) variations across different anode and cathode placements. **a.** MT for each spinal root (top) and each muscle (bottom) across the different anode placements in (Fig. 5a). **b.** MT for each spinal root (top) and each muscle (bottom) across the different cathode placements in (Fig. 5j). MTs were clipped at 65 a.u. for better visualization of the differences across paradigms. Abbreviations: arbitrary units (a.u.), extensor carpi radialis (ECR), flexor capri radialis (FCR), abductor pollicis brevis (APB), interosseous (Int.).

