## Supplementary Information for "Virtual prototyping of non-invasive spinal cord electrical stimulation targeting upper limb motor function"

| Segment | Reported avg. segment length[1] (mm) | Reported avg. REZ length[1] (mm) | Reported avg. segment length percentage[2] (%) | Calculated segment length (mm) | Calculated ratio (avg. REZ length / avg. segment length) | Calculated REZ length (mm) |
|---|---|---|---|---|---|---|
| **C1** | 12.60* | 11.30* | 1.60 | 7.57 | 0.90 | 6.79 |
| **C2** | 12.60 | 11.30 | 2.20 | 10.41 | 0.90 | 9.34 |
| **C3** | 13.30 | 9.40 | 3.50 | 16.57 | 0.71 | 11.71 |
| **C4** | 13.90 | 11.60 | 3.50 | 16.57 | 0.83 | 13.83 |
| **C5** | 15.00 | 12.40 | 3.50 | 16.57 | 0.83 | 13.70 |
| **C6** | 13.60 | 11.50 | 3.30 | 15.62 | 0.85 | 13.21 |
| **C7** | 11.70 | 11.40 | 3.20 | 15.15 | 0.97 | 14.76 |
| **C8** | 12.50 | 11.80 | 3.40 | 16.09 | 0.94 | 15.19 |
| **T1** | 13.00 | 12.00 | 3.60 | 17.04 | 0.92 | 15.73 |
| **T2** | 16.40 | 13.10 | 3.90 | 18.46 | 0.80 | 14.75 |
| **T3** | 16.10 | 15.30 | 4.40 | 20.83 | 0.95 | 19.79 |
| **T4** | 22.80 | 18.40 | 5.00 | 23.67 | 0.81 | 19.10 |
| **T5** | 22.10 | 19.60 | 5.10 | 24.14 | 0.89 | 21.41 |
| **T6** | 24.80 | 18.80 | 5.60 | 26.51 | 0.76 | 20.09 |
| **T7** | 24.50 | 20.10 | 5.60 | 26.51 | 0.82 | 21.75 |
| **T8** | 25.50 | 19.00 | 5.40 | 25.56 | 0.75 | 19.04 |
| **T9** | 23.90 | 20.40 | 5.10 | 24.14 | 0.85 | 20.60 |
| **T10** | 25.30 | 19.30 | 4.70 | 22.25 | 0.76 | 16.97 |
| **T11** | 22.40 | 15.50 | 4.30 | 20.35 | 0.69 | 14.08 |
| **T12** | 18.40 | 15.60 | 3.90 | 18.46 | 0.85 | 15.65 |
| **L1** | 17.50 | 15.20 | 3.60 | 17.04 | 0.87 | 14.80 |
| **L2** | 14.00 | 11.80 | 2.80 | 13.25 | 0.84 | 11.17 |
| **L3** | 11.20 | 11.20 | 2.40 | 11.36 | 1.00 | 11.36 |
| **L4** | 12.50 | 11.60 | 2.20 | 10.41 | 0.93 | 9.66 |
| **L5** | 9.70 | 11.20 | 1.70 | 8.05 | 1.15 | 9.29 |
| **S1** | 9.70* | 11.20* | 1.50 | 7.10 | 1.15 | 8.20 |
| **S2** | 9.70* | 11.20* | 1.60 | 7.57 | 1.15 | 8.74 |
| **S3** | 9.70* | 11.20* | 1.40 | 6.63 | 1.15 | 7.65 |
| **S4** | 9.70* | 11.20* | 1.30 | 6.15 | 1.15 | 7.10 |

**Supplementary Table 1.** Reported[1] and calculated segment-specific morphometric parameters used for the 3D rootlets model generation. *These parameters were not reported in the referenced publication but were extrapolated from parameters of the closest reported segments. Abbreviations: average (avg.), millimeter (mm), root entry zone (REZ).

| Segment | Reported avg. number of rootlets[1] | | Calculated number of rootlets | | Reported avg. root diameter[1] (mm) | | Calculated rootlet diameter (mm) | |
|---|---|---|---|---|---|---|---|---|
| | P | A | P | A | P | A | P | A |
| **C1** | 8.33* | 8.33* | 5 | 5 | 4.34* | 2.79* | 0.87 | 0.56 |
| **C2** | 8.33 | 8.33 | 7 | 7 | 4.34 | 2.79 | 0.62 | 0.40 |
| **C3** | 8.25 | 6.25 | 11 | 8 | 4.55 | 2.63 | 0.41 | 0.33 |
| **C4** | 8.87 | 7.62 | 11 | 9 | 4.84 | 2.85 | 0.44 | 0.32 |
| **C5** | 9.22 | 8.14 | 10 | 9 | 5.43 | 3.78 | 0.54 | 0.42 |
| **C6** | 8.77 | 6.85 | 10 | 8 | 5.08 | 3.29 | 0.51 | 0.41 |
| **C7** | 7.77 | 7.25 | 10 | 9 | 5.50 | 2.92 | 0.55 | 0.32 |
| **C8** | 7.66 | 6.62 | 10 | 9 | 5.18 | 2.19 | 0.52 | 0.24 |
| **T1** | 6.00 | 4.66 | 8 | 6 | 3.85 | 1.80 | 0.48 | 0.30 |
| **T2** | 5.00 | 4.33 | 6 | 5 | 3.35 | 1.82 | 0.56 | 0.36 |
| **T3** | 4.88 | 4.22 | 6 | 6 | 3.37 | 1.33 | 0.56 | 0.22 |
| **T4** | 4.44 | 4.33 | 5 | 4 | 3.21 | 1.38 | 0.64 | 0.34 |
| **T5** | 4.00 | 4.88 | 4 | 5 | 3.29 | 1.47 | 0.82 | 0.29 |
| **T6** | 4.55 | 4.44 | 5 | 5 | 3.62 | 1.71 | 0.72 | 0.34 |
| **T7** | 4.22 | 4.66 | 5 | 5 | 3.27 | 1.71 | 0.65 | 0.34 |
| **T8** | 4.55 | 4.66 | 5 | 5 | 3.73 | 1.64 | 0.75 | 0.33 |
| **T9** | 5.44 | 4.66 | 6 | 5 | 3.83 | 1.43 | 0.64 | 0.29 |
| **T10** | 4.88 | 4.55 | 4 | 4 | 3.71 | 1.67 | 0.93 | 0.42 |
| **T11** | 5.33 | 3.88 | 5 | 4 | 3.20 | 1.32 | 0.64 | 0.33 |
| **T12** | 6.22 | 4.11 | 6 | 4 | 3.53 | 1.47 | 0.59 | 0.37 |
| **L1** | 7.22 | 5.33 | 7 | 5 | 3.84 | 2.45 | 0.55 | 0.49 |
| **L2** | 8.00 | 5.88 | 8 | 6 | 4.60 | 2.94 | 0.57 | 0.49 |
| **L3** | 8.55 | 6.33 | 8 | 6 | 4.26 | 2.70 | 0.53 | 0.45 |
| **L4** | 7.55 | 5.00 | 6 | 4 | 4.40 | 3.19 | 0.73 | 0.80 |
| **L5** | 8.22 | 5.33 | 7 | 5 | 4.98 | 2.83 | 0.71 | 0.57 |
| **S1** | 8.22* | 5.33* | 6 | 4 | 4.98* | 2.83* | 0.83 | 0.71 |
| **S2** | 8.22* | 5.33* | 7 | 4 | 4.98* | 2.83* | 0.71 | 0.71 |
| **S3** | 8.22* | 5.33* | 5 | 3 | 4.98* | 2.83* | 1.00 | 0.94 |
| **S4** | 8.22* | 5.33* | 5 | 3 | 4.98* | 2.83* | 1.00 | 0.94 |

**Supplementary Table 2.** Reported[1] and calculated segment- and position-specific morphometric parameters used for the 3D rootlets model generation. *These parameters were not reported in the referenced publication but were extrapolated from parameters of the closest reported segments. Abbreviations: average (avg.), millimeter (mm), posterior (P), anterior (A)

| Root pairs | Left | | Right | |
|---|---|---|---|---|
| | Reported occurrences[1] (n=9) | Modelled | Reported occurrences[1] (n=9) | Modelled |
| C2-C3 | 7 | ✓ | 4 | X |
| C3-C4 | 9 | ✓ | 6 | ✓ |
| C4-C5 | 6 | ✓ | 4 | X |
| C5-C6 | 7 | ✓ | 7 | ✓ |
| C6-C7 | 7 | ✓ | 5 | ✓ |
| C7-C8 | 6 | ✓ | 5 | ✓ |
| C8-T1 | 2 | X | 2 | X |
| L1-L2 | 2 | X | 1 | X |
| L2-L3 | 0 | X | 0 | X |
| L3-L4 | 2 | X | 1 | X |
| L4-L5 | 1 | X | 0 | X |
| L-S1 | 1 | X | 0 | X |

**Supplementary Table 3.** Reported occurrences of inter-root anastomoses[1] and their inclusion in the modeling of the 3D structures in our model. Only anastomoses with occurrences 5 or more occurrences (>50%) were considered in the modeling.

| Tissue | Maximum step (X, Y, Z in mm) | Geometry resolution (X, Y, Z in mm) |
|---|---|---|
| Rootlets | 0.22, 0.22, 0.22 | 0.18, 0.18, 0.18 |
| Peripheral nerves | 2.0, 2.0, 2.0 | 1.0, 1.0, 1.0 |
| Cranial nerve branches (including vagus nerve) | 2.0, 2.0, 2.0 | 1.45, 1.45, 1.45 |
| Gray matter, white matter (SC), cerebrospinal fluid and dura mater | 1.0, 1.0, 2.0 | 1.0, 1.0, 1.0 |
| Skin | 0.5, 0.5, 1.0 | 0.5, 0.5, 1.0 |
| Tissue hole filler | 3.0, 3.0, 5.0 | 3.0, 3.0, 10.0 |
| Other tissues | 2.0, 2.0, 10.0 | 2.0, 2.0, 2.0 |

**Supplementary Table 4.** Voxelization parameters used for the different tissues in the 3D whole-body model. Abbreviations: millimeter (mm), spinal cord (SC).

### Supplementary methods:

#### End-nodal recruitment:

We noticed a phenomenon of end-nodal recruitment for some axons, a known limitation of conductance-based compartmental cable models[3,4]. To ensure the interpretability of our simulation results, we sought to avoid such activations. Therefore, we introduced two methods of avoiding phantom end-node activations and assessed their ability in shifting the initiation sited of action potentials (ISAPs), in addition to their effect on the threshold of activation (**Supplementary Fig. 1**). Based on our results, we replaced terminal nodes with passive nodes to avoid end-node recruitment[3] (**Supplementary Fig. 1**).

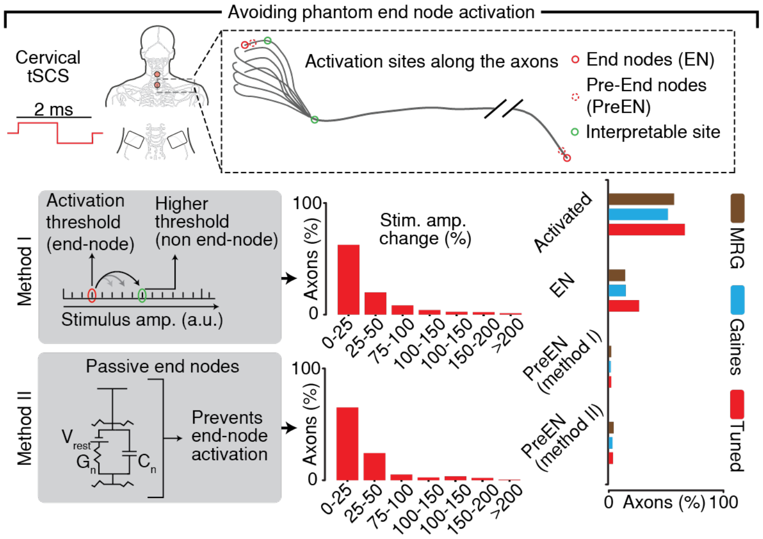


**Supplementary Fig. 1.** Proposed methods for avoiding non-interpretable end-node activations observed in simulations of extracellularly stimulating conductance-based compartmental cable models[3,4]. Top: electrode configuration where end-nodes activations could occur and end-nodes highlighted along a sketch of nerve axons. Bottom: Proposed methods for avoiding phantom end-node activations. Method I: Avoiding end-node activation by increasing stimulus amplitude until a new activation site is present. Method II: Avoiding end-node activation by replacing the nodes of Ranvier at the termini with passive nodes[3], preventing any activation to be initiated from them. The red bar plots show histograms of the relative stimulus threshold changes caused by using each of the methods on the tuned cable model, while the vertical bar chart shows the percentage of axons exhibiting end-node activation as well as the percentage of axons where the activation moved to only the next node after applying each of the two methods using each of the three different cable models. Abbreviations: millisecond (ms), end nodes (EN), pre-end nodes (PreEN), nodal conductance ($G_n$), nodal capacitance ($C_n$), resting membrane potential ($V_{rest}$).

#### Axon model tuning:

As mentioned in the methods' subsection "Axon model tuning", we constructed a parameter space composed of all parameters, for which we found reported variations[5–14] of ion channel properties between somatosensory afferents and motor efferents (**Supplementary Table 4**). We ensured that all sampled parameter combinations preserved the reported property differences[5–14] (**Supplementary Table 4**).

| Ion channel property | Sensory / motor difference |
| --- | --- |
| Persistent sodium conductance ($gNa_p$) | Sensory > motor[5–7] |
| Slow potassium conductance ($gK_s$) | Sensory > motor (ratio = 1.55)[8,9] |
| Fast potassium conductance ($gK_f$) | Sensory < motor (ratio = 0.6)[10] |
| Leakage conductance (gL) | Sensory > motor (ratio = 1.5)[11] |
| Persistent sodium speed ($\alpha m_pA$, $\beta m_pA$) | Sensory < motor[12] |
| Fast sodium speed (αmA, βmA) | Sensory < motor[12] |
| Sodium inactivation speed (αhA, βhA) | Sensory > motor[8,12] |
| Slow potassium speed (αsA, βsA) | Sensory > motor[13] |
| Slow potassium half-activation voltage (αsB, βsB) | Sensory < motor[10,14] |
| Fast potassium speed (αnA, βnA) | Sensory > motor[13] |

**Supplementary Table 5.** Ion channel differences between sensory and motor nerve fibers as reported in the literature across different species. α parameters refer to opening of ion channels, while β parameters refer to closing of ion channels.

| | | Node | | | FLUT | | MYSA, STIN |
|---|---|---|---|---|---|---|---|
| **Voltage and time dependent parameters** | A($ms^{-1}$) | B(mV) | C(mV) | A($ms^{-1}$) | B(mV) | C(mV) | |
| αm | 1.86 | 21.4 | 10.3 | – | – | – | - |
| βm | 0.086 | 25.7 | 9.16 | – | – | – | - |
| αmp | 0.01 | 27 | 10.2 | – | – | – | - |
| βmp | 0.00025 | 34 | 10 | – | – | – | - |
| αh | **0.0728** | 114.0 | 11.0 | – | – | – | - |
| βh | 2.3 | 31.8 | 13.4 | – | – | – | – |
| αn | - | - | - | 0.0462 | -83.2 | 1.1 | - |
| βn | - | - | - | 0.0824 | **-60.8684** | 10.5 | - |
| αs | 0.3 | -27** | -5 | - | - | - | - |
| βs | **0.048** | 10 | -1 | - | - | - | - |
| **Channel Conductances** | | (mS cm-2) | | | (mS cm-2) | | (mS cm-2) |
| Persistent Sodium | | 10 | | | – | | - |
| Fast Sodium | | 3000 | | | – | | - |
| Slow Potassium | | 80 | | | - | | - |
| Fast Potassium | | - | | | 20 | | - |
| Leak | | 7 | | | - | | - |
| Passive | | - | | | 0.1* | | 1 (MYSA), 0.1 (STIN)* |
| **Reversal Potentials** | | (mV) | | | (mV) | | |
| Sodium | | 50 | | | – | | - |
| Slow Potassium | | -90.0 | | | - | | - |
| Fast Potassium | | - | | | -90.0 | | - |
| Leak | | -90.0 | | | - | | - |
| **Resting potential (mV)** | | | | -78.05 | | | |

**Supplementary Table 6.** Parameters used in the newly-tuned MRG-type model for somatosensory afferent axons. Highlighted are values that deviated from those reported in the original MRG-model[8]. *This parameter scales with the ratio of segment diameter to fiber diameter as in the original MRG-model[8]. **To avoid spontaneous activation, this parameter was modified for very small fibers (diameter < 5.16μm) but maintained sensory/motor difference (23 mV). Abbreviations: fluted (FLUT), stereotyped Internode (STIN), myelin sheath attachment (MYSA), ion channel conductance (g), ion channel opening rate (α), ion channel closing rate (β), ion channel maximum speed of transition (A), ion channel half-activation voltage (B), ion channel response slope (C), sodium activation (m), persistent sodium activation (mp), sodium inactivation (h), fast potassium activation (n), slow potassium activation (s), millisecond (ms), millivolt (mV), milli Siemens (mS), centimeter (cm)

| | Node | | | FLUT | | | MYSA, STIN |
|---|---|---|---|---|---|---|---|
| **Voltage and time dependent parameters** | A($ms^{-1}$) | B(mV) | C(mV) | A($ms^{-1}$) | B(mV) | C(mV) | |
| αm | **1.95789** | 21.4 | 10.3 | – | – | – | - |
| βm | **0.090526** | 25.7 | 9.16 | – | – | – | - |
| αmp | **0.016** | 27 | 10.2 | – | – | – | - |
| βmp | **0.00036959** | 34 | 10 | – | – | – | - |
| αh | **0.0728** | 114.0 | 11.0 | – | – | – | - |
| βh | **2.1789** | 31.8 | 13.4 | – | – | – | – |
| αn | - | - | - | **0.037149** | -83.2 | 1.1 | - |
| βn | - | - | - | **0.0515** | -66 | 10.5 | - |
| αs | 0.3 | -27 | -5 | - | - | - | - |
| βs | 0.03 | 10 | -1 | - | - | - | - |
| **Channel Conductances** | (mS cm-2) | | | (mS cm-2) | | | (mS cm-2) |
| Persistent Sodium | 10 | | | – | | | - |
| Fast Sodium | 3000 | | | – | | | - |
| Slow Potassium | 80 | | | - | | | - |
| Fast Potassium | - | | | 20 | | | - |
| Leak | 7 | | | - | | | - |
| Passive | - | | | 0.1* | | | 1 (MYSA), 0.1 (STIN)* |
| **Reversal Potentials** | (mV) | | | (mV) | | | |
| Sodium | 50 | | | – | | | - |
| Slow Potassium | -90.0 | | | - | | | - |
| Fast Potassium | - | | | -90.0 | | | - |
| Leak | -90.0 | | | - | | | - |
| **Resting potential (mV)** | -79.5 | | | | | | |

**Supplementary Table 7.** Parameters used in the tuned MRG-type model for motor efferent axons[8]. Highlighted are values that deviated from those reported in the original MRG-model. *This parameter scales with the ratio of node diameter to fiber diameter as in the original MRG-model[8]. Abbreviations: fluted (FLUT), stereotyped Internode (STIN), myelin sheath attachment (MYSA), ion channel conductance (g), ion channel opening rate (α), ion channel closing rate (β), ion channel maximum speed of transition (A), ion channel half-activation voltage (B), ion channel response slope (C), sodium activation (m), persistent sodium activation (mp), sodium inactivation (h), fast potassium activation (n), slow potassium activation (s), millisecond (ms), millivolt (mV), milli Siemens (mS), centimeter (cm)
